# Engineered Living Glues for Autonomous Detection and On-Demand Treatment of Inflammatory Bowel Disease

**DOI:** 10.1101/2025.01.04.630840

**Authors:** Changhao Ge, Shanshan Jiang, Xiaomin Dong, Xiaoyu Jiang, Weiliang Zhi, Yunqing Xiang, Peilang Yang, Qian Zhang, Xin Chen, Yan Liu, Shuqiang Huang, Yifan Liu, Jing Lin, Bolin An, Peng Huang, Chao Zhong

## Abstract

Smart biomaterials capable of autonomously sensing pathological conditions and executing targeted medical interventions offer therapeutic advantages over conventional passive biomaterials. However, their development remains a considerable challenge. Here, we introduce a therapeutic “living glue” for the automatic detection and on-demand treatment of inflammatory bowel disease (IBD). This living glue employs a genetically engineered, non-pathogenic *Escherichia coli* with a highly sensitive blood-inducible gene circuit to monitor gastrointestinal bleeding, a key indicator of severe IBD. Upon detection, the bacteria respond by producing potent adhesive and therapeutic proteins around bleeding sites, enabling robust attachment to inflamed tissues and sustained treatment. In a dextran sulfate sodium (DSS)-induced mouse model, a single rectal administration of the living glue markedly improved weight recovery, reversed colonic shortening, and reduced intestinal bleeding. Additionally, the living glue decreased intestinal inflammation, promoted mucosal repair, and restored gut barrier integrity, demonstrating comprehensive therapeutic effects in alleviating IBD symptoms. This study highlights the potential of integrating programmable, living components into biomaterials for autonomous, targeted, and enhanced medical interventions.

## Introduction

In the rapidly evolving field of biomedical science, biomaterials are indispensable; however, they often remain limited by their passive and non-responsive nature^1–3^. Traditional biomaterials typically require external intervention to realize their full therapeutic potential, as they inherently lack the ability to sense and adapt to complex pathological conditions^4^. For example, hydrogel-based adhesives have shown promising efficacy in tissue adhesion, hemostasis, and wound healing^5, 6^. However, their use within the body generally depends on sophisticated diagnostic devices to locate bleeding sites, often requiring invasive surgical procedures. This reliance is especially problematic for conditions like gastrointestinal bleeding, which is difficult to detect and treat precisely with “static” biomaterials^7^. In contrast, living organisms can produce biomaterials that dynamically adapt to their surroundings and respond on demand with high programmability^8^. Marine barnacles, for instance, secrete cement proteins to firmly anchor themselves to underwater surfaces in preferred habitats, while sandcastle worms selectively apply glue proteins to assemble their nests^9^. Inspired by such natural examples of programmed dynamic adhesion, interest has grown in biomaterials with similar “programmability” to address unmet medical needs^3, 10^. Notable advancements include the development of an orally administered, glucose-responsive micelle material for type 1 diabetes management^11^ and a thrombin-responsive DNA nanodevice for precise thrombosis treatment^12^. Despite recent progress in biomaterial innovation, the design of intelligent, responsive adhesives that autonomously target disease sites and execute therapeutic actions in response to specific biological cues or user inputs remains a daunting challenge^13^.

Synthetic biology, which aims to program biological systems for user-defined functions^14, 15^, has merged with materials science to create environmentally responsive biomaterials with “living” characteristics, often referred to as engineered living materials (ELMs)^16, 17^. ELMs harness tailored cellular functions to produce versatile biomaterials that mimic biological systems’ dynamic and autonomous behaviors^16, 18, 19^. This emerging field offers a new paradigm in material synthesis and applications, with transformative potential for intelligent materials and cellular devices^17, 20^. For example, synthetic biology tools allow researchers to program microbes to precisely regulate material production and assembly, facilitating their *in situ* application. Examples include engineering *Bacillus subtilis* to secrete recombinant extracellular polymeric matrix for enhanced biocatalysis^21, 22^; cultivating self-pigmenting, sustainable textiles using genetically modified cellulose-producing bacteria^23^; and incorporating environmentally responsive circuits into *Escherichia coli* adhesive biofilms for site-specific mechanical functions^24,25^. Although recent progress has been made in developing ELMs, their application in medical contexts, particularly for autonomously detecting and treating hard-to-reach diseases, remains in early exploratory stages, highlighting a crucial area for future research and development^2, 17, 20^.

Here, we present a therapeutic living glue for the on-demand detection and continuous treatment of gastrointestinal disorders, with a focus on inflammatory bowel diseases (IBD). Our strategy harnesses genetically modified *E. coli* equipped with a highly sensitive blood-inducible gene circuit to trigger the localized production of both adhesive and therapeutic proteins. This design enables the engineered bacteria to detect and respond to bleeding, a hallmark of severe IBD episodes caused by compromised gut barriers^26^, allowing precise recognition and robust adherence to affected sites for sustained treatment (**Fig. 1**). Specifically, our approach integrates three key features. First, we constructed a high-performance blood-sensing gene circuit incorporating a multi-layered transcriptional amplifier and optimizing promoters to enhance responsiveness, enabling the engineered microbes to precisely monitor gastrointestinal bleeding with a fluorescence signal amplification exceeding 100-fold at blood concentrations as low as 100 ppm. Second, we incorporated barnacle-derived cement proteins, such as CP19K and CP43K^27, 28^, as biocompatible adhesive components. Upon detecting blood, the engineered microbes produce and secrete recombinant cement proteins, creating anti-inflammatory adhesive matrices that protect inflamed tissues. Third, recognizing that mucosal barrier restoration is essential in managing IBD, the bacteria are engineered to secrete TFF3, an 11 kDa peptide essential for mucosal healing and rapid intestinal wound repair^29^, in response to blood. This therapeutic glue matrix on wound tissue not only serves as a protective barrier against further intestinal inflammation but also retains therapeutic bacteria for *in situ* disease resolution (**Fig. 1**). In DSS-induced IBD mouse models, the living glue demonstrated accurate blood signal detection, durable adhesion, reduced bleeding, alleviated inflammation, and enhanced intestinal barrier repair. This study underscores the remarkable therapeutic potential of living materials with programmable functionalities, offering intelligent strategies for treating complex medical conditions.

**Figure 1:**
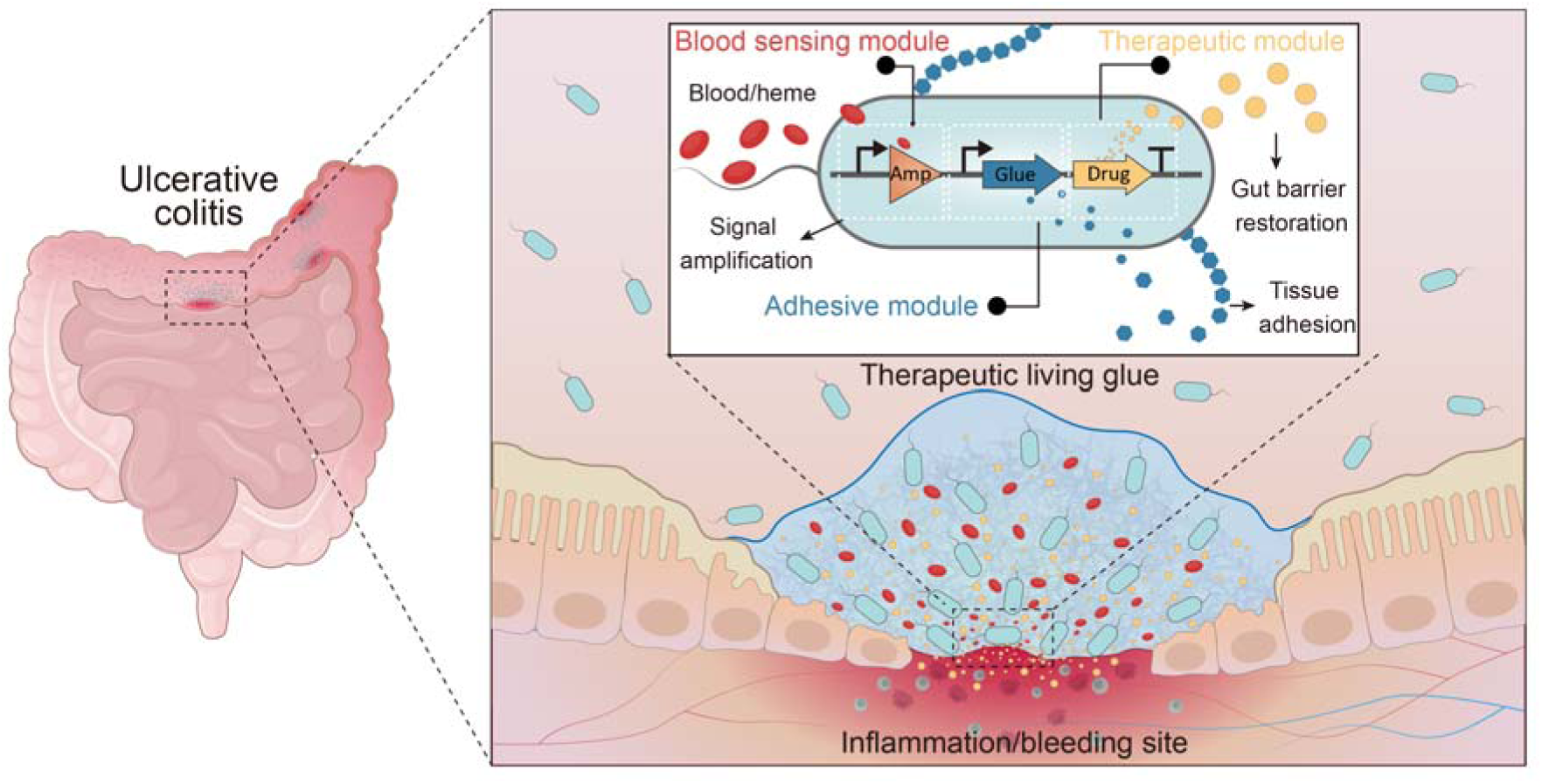
Schematic of the therapeutic living glue system for precise targeting of bleeding sites and prolonged treatment of IBD. An artificial genetic circuit comprising a blood-sensing module, an adhesive module, and a therapeutic module is incorporated into engineered *E. coli* to achieve multiple functions. The blood-sensing module, including a blood-inducible promoter and a transcriptional amplifier, allows sensitive detection of the bleeding symptom associated with ulcerative colitis. The adhesive and therapeutic modules are designed to produce potent adhesive and therapeutic protein elements respectively, enabling targeted tissue adherence and gut barrier restoration.

## Results

### A Blood-Inducible Gene Circuit with High Signal-to-Noise Ratio

To use blood as a biomarker for intelligent therapeutic living materials in IBD, we redesigned a previously reported blood/heme-responsive genetic circuit^30^. This circuit consists of a heme transporter (ChuA), a heme-responsive transcriptional repressor (HrtR), and a heme-specific promoter (P*_L_*_(*HrtO*)_) (**Fig. 2a, Supplementary Fig. 1a**). Initially designed for gut bleeding detection in an ingestible, ultrasensitive electronic device, this system used genetically modified bacteria to monitor gut blood signals in real-time^31^. However, while the signal-to-noise ratio of this circuit was effective for blood detection using luciferase and luminance as readouts, it produced insufficient bioactive molecules for therapeutic applications (**Supplementary Fig. 1b**). To enhance sensitivity and enable therapeutic use, we integrated a multi-layered transcriptional amplifier to bridge the biosensing and output modules (**Fig. 2a**). Transcriptional amplifiers, characterized by positive feedback loops and sequential signaling cascades, relay signals to stronger promoters and increase cellular response to specific environmental cues^31^. We used a three-layer amplifier cascade with transcription factors and promoters, each stage sequentially activating the next, significantly boosting the final output^32^ (**Fig. 2a, 2b**).

**Figure 2:**
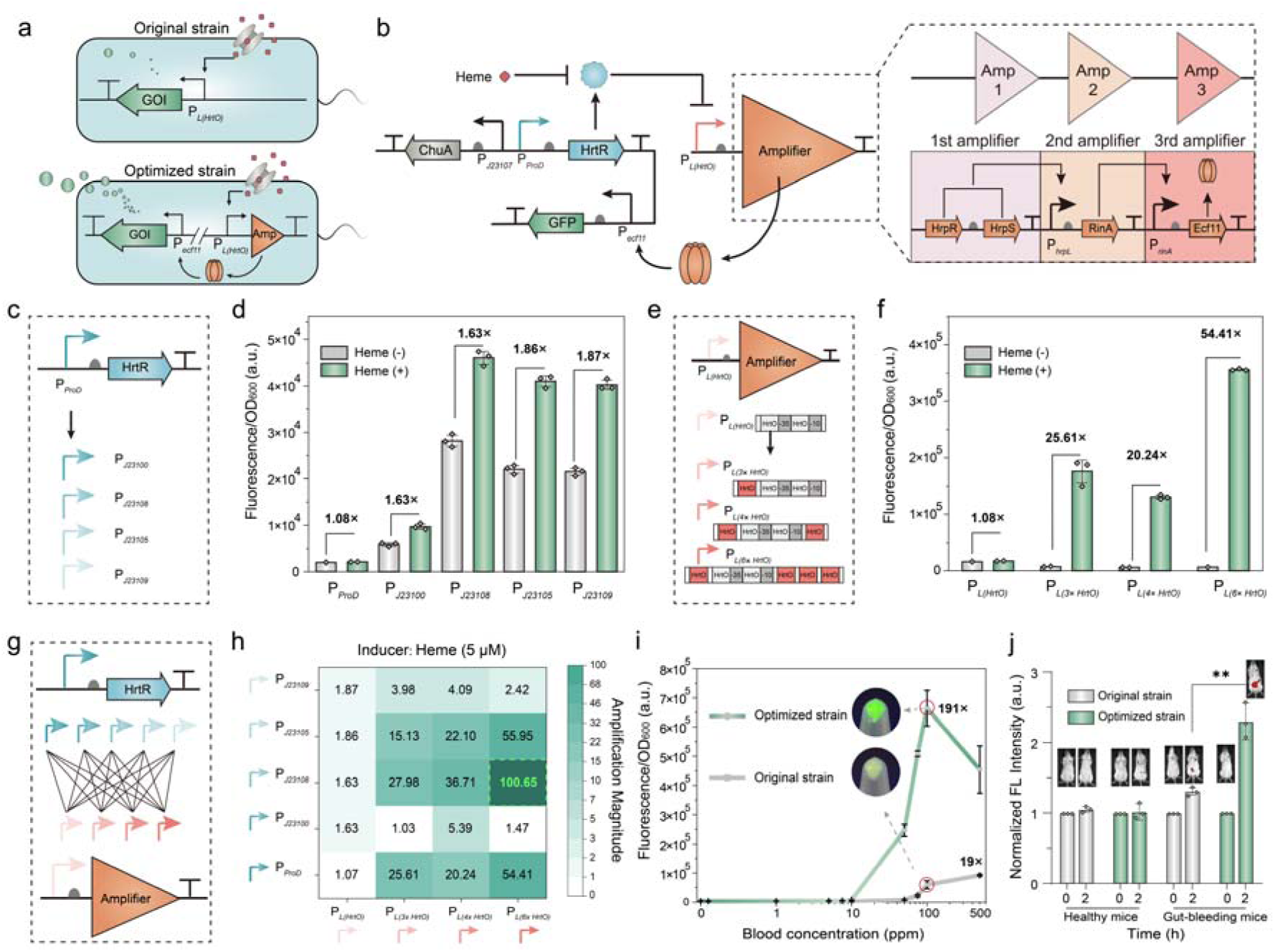
Construction of a blood-inducible gene circuit with high signal-to-noise ratios. **(a)** Diagram illustrating the gene constructs in the original and optimized strains. **(b)** Schematic of the optimized blood-inducible gene circuit, which incorporates a transcriptional amplifier bridging the blood-sensing module with the output adhesive and therapeutic module. **(c)** Illustration of the constitutive promoters chosen to drive HrtR repressor expression. (**d)** Comparative analysis of fluorescent intensity in engineered strains using various constitutive promoters for HrtR expression (*n* = 3 independent samples, mean[±[standard deviation (SD)). The heme inducer concentration was fixed at 5 μM. **(e)** Diagram of blood-responsive promoter variants. **(f)** Comparison of fluorescent intensity in engineered strains with different blood-responsive promoter variants (*n* = 3 independent experiments, mean ± SD). In this setup, the promoter driving HrtR expression was P*_ProD_*, with the heme inducer concentration held at 5 μM. **(g)** Schematic for library construction of genetic circuits combining two types of promoters. **(h)** Heat map showing blood-inducible performance across 20 constructed strains (*n* = 3 independent experiments, mean ± SD) at heme inducer concentration of 5 μM. Increasing green color intensity corresponds to higher induction of the gene circuits. **(i)** Comparative dose-response analysis between the optimized and prototype strains at varying fresh mouse blood concentrations (*n* = 3 independent experiments, mean ± SD), including fluorescent images of the *E. coli* negative control, prototype control, and optimized strain. **(j)** Semi-quantitative analysis of blood-induced miRFP680 expression in bacteria within mice 2-hour post-rectal injection (*n* =[3 independent experiments, mean ± SD), with representative images from *in vivo* fluorescence imaging (FLI). Excitation wavelength for miRFP680 = 640 nm; emission wavelength = 680 nm.

To test this amplifier’s efficacy, we integrated the amplifier-expressing cassette, regulated by the heme-responsive promoter P*_L_*_(*HrtO*)_, into plasmids and transformed them into bacteria expressing ChuA, HrtR, and a GFP reporter regulated by the third-layer amplifier promoter P*_ecf11_*(**Fig. 2b, Supplementary Fig. 1c**). The results showed that low HrtR levels led to robust GFP expression, significantly higher than the original strain (**Supplementary Fig. 1d**). However, GFP expression remained high even without an inducer, likely due to insufficient HrtR failing to repress P*_L_*_(*HrtO*)_ promoters. In contrast, when HrtR expression was increased using a high-copy plasmid, GFP levels dropped sharply, likely due to excessive repression (**Supplementary Fig. 1d**). These findings highlight the need to balance HrtR expression for optimal circuit function.

To achieve this balance, we paired a low-copy plasmid encoding the amplifier with a high-copy plasmid harboring for HrtR expression. We adjusted HrtR expression by replacing its intrinsic promoter, P*_proD_*, with progressively weaker Anderson promoters (P*_J23100_*, P*_J23108_*, P*_J23105_*, and P*_J23109_*) (**Fig. 2c)**. These changes increased GFP expression by 1.5-to 2.0-fold, with weaker promoters notably boosting sensitivity (**Fig. 2d)**. However, these modifications also led to high GFP expression leakage without inducers (**Fig. 2d)**. To reduce this leakage, we modified the blood-responsive promoter by adding more copies of HrtO, the operon sequence of P*_L_*_(*HrtO*)_. We hypothesized that more HrtO sites would increase HrtR’s binding affinity to the promoter, reducing promoter activity in the uninduced state. Configurations with P*_L_*_(*3×*_ *_HrtO_*_)_, P*_L_*_(*4×*_ *_HrtO_*_)_, and P*_L_*_(*6×*_ *_HrtO_*_)_ were tested (**Fig. 2e)**. The P*_L_*_(*6×*_ *_HrtO_*_)_ variant produced a 54-fold increase in induction when used in the blood-inducible circuit (**Fig. 2f)**, indicating the importance of modulating the transcriptional repression of HrtR to optimize the blood-responsive sensor performance.

We then combined these optimized promoter configurations to create 20 novel blood-inducible gene circuits (**Fig. 2g**). Each circuit was evaluated in *E. coli* with a 5 µM heme inducer, and heatmap analysis identified the configuration with the greatest induction (100.6-fold), which used promoter P*_J23108_*to drive HrtR expression and P*_L_*_(*6×*_ *_HrtO_*_)_ to activate the three-layer amplifier (**Fig. 2h**). We subsequently tested the circuit’s responsiveness across a range of heme and blood concentrations. The optimized circuit achieved a 108-fold induction with 10 µM heme and a 191-fold induction using 100 ppm of fresh mouse blood (**Fig. 2i, Supplementary Fig. 2, 3**), a significant improvement over the original circuit’s 10 to 19-fold increase in gene expression (**Fig. 2i)**.

Next, we tested the *in vivo* performance of the optimized blood-responsive circuit by driving the expression of miRFP680, a near-infrared (NIR) light-excited fluorescent protein activated by heme cofactors, enabling real-time *in vivo* imaging^33^. We administered engineered *E. coli* strains containing either the prototype or optimized circuits to mouse models, both with and without induced gut bleeding. Within two hours, significant fluorescence signals were observed only in the IBD gut-bleeding mice that received the strain with the optimized circuit (**Fig. 2j**). This result confirmed that our optimized blood-responsive circuit enabled engineered microbes to detect blood molecules and produce substantial amounts of fluorescent protein *in vivo*, showing promise for scalable production of therapeutic agents for treating bleeding-related disorders.

### Design and Characterization of Blood-Responsive Living Glues

We proposed a probiotic-incorporated living glue to treat gastrointestinal diseases by adhering to intestinal wounds for extended treatment efficacy. Previous efforts involved engineering *E. coli* curli biofilm (composed of CsgA protein) with mussel foot proteins to create biofilm-enabled adhesives^24, 25^. However, concerns regarding CsgA’s inflammatory potential and association with pathogenicity^34, 35^ have led us to explore alternative designs with safer profiles. The natural adhesive properties of barnacle cement proteins (CPs), particularly CP43K and CP19K, are known to facilitate self-assembly into amyloid-like fibers under aqueous conditions^36^ (**Fig. 3a**). Meanwhile, *in vitro* experiments showed that purified CPs possess moderate anti-inflammatory effects^37^ (**Supplementary Fig. 4a**). We thus reasoned that engineered microbes secreting these CPs could create extracellular adhesive matrices around bacterial cells, forming an anti-inflammatory living glue that robustly adheres to inflamed intestinal tissues. Cytotoxicity tests also indicated that barnacle CPs have high biocompatibility (**Supplementary Fig. 4b**), indicating their potential for disease treatment.

**Figure 3:**
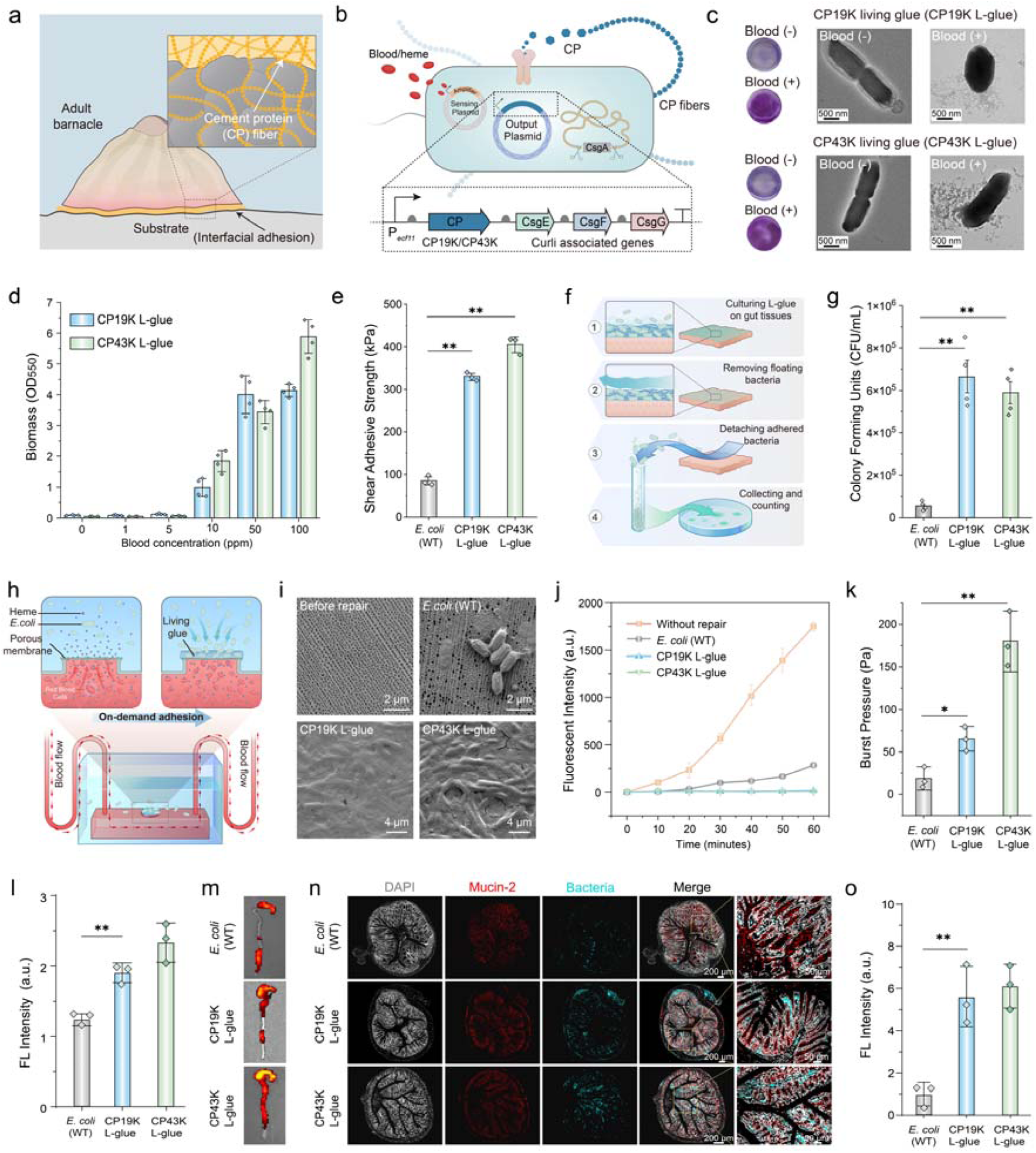
Design and characterization of the engineered living glue materials. **(a)** Diagram illustrating the barnacle and its adhesive matrix composed of cement protein fibers. **(b)** Genetic circuit design for engineered living glue materials. **(c)** CV staining and TEM images of CP19K and CP43K L-glues. **(d)** Production of CP19K and CP43K L-glues as a function of blood concentration (*n□*=*□*4 independent experiments, mean ± SD). **(e)** Lap-shear test schematic (left) and shear adhesive strength measurements for wild-type *E. coli* control, CP19K L-glue, and CP43K L-glue (*n □* =*□*3 independent experiments, mean ±*□*SD). (**f)** Diagram illustrating *in vitro* adhesion procedures for testing living glue on intestinal tissues. **(g)** Quantification of bacteria maintained on intestinal tissues post-washing by CFU (*n* = 4 independent experiments, mean ± SD). (**h**) Diagram illustrating the targeted, on-demand adhesion of engineered L-glue on a damaged microfluidic setup. A 1.2 mm diameter hole was intentionally introduced into the microfluidic channel and covered with a porous aluminum membrane. The membrane’s 200 nm pores prevented bacterial cells from entering the channel while allowing the diffusion of heme molecules, which subsequently induced surrounding bacteria to produce protein-based adhesive components. (**i**) SEM images showing the porous aluminum membranes covering the microfluidic channel before and after engineered microbial repair. (**j**) Fluorescence leakage assays for evaluating the repair effects (n = 3 independent experiments, mean ± SD). A 50 mM Cy3 solution was pumped through the device at a flow rate of 250 mL/h, and fluorescence in the medium adjacent to the leaking membrane was monitored every 10 minutes. (**k**) Comparison of burst pressure resistance between wild-type *E. coli* and two engineered living glues (n = 3 independent experiments, mean ± SD). **(l)** Semi-quantitative analysis of *ex vivo* colon tissues 24 hours post-rectal injection of engineered bacteria in IBD mice (*n* = 3 independent experiments, mean ± SD). **(m)** Representative fluorescent images of *ex vivo* colon tissues 24 hours post-rectal injection of engineered bacteria in IBD mice **(n)** Fluorescence In Situ Hybridization (FISH) and **(o)** Semi-quantitative analysis of *ex vivo* colon tissues 24 hours post-rectal injection of engineered bacteria in IBD mice (*n* = 3 independent experiments, mean ± SD). Staining details: Grey for DAPI, Red for Mucin-2, Cyan for *E. coli* K-12 MG1655 by 16S rRNA FISH, and Black for overlay. Scale bars: 200 and 50 μm, respectively.

We employed the *E. coli* curli system’s secretion machinery to secrete recombinant adhesive proteins fused with the CsgA signal peptide and either CP43K or CP19K at the C-terminus. Protein expression was driven by our optimized blood-inducible circuit (**Fig. 3b**). Co-expression of curli-associated proteins CsgE, CsgF, and CsgG improved secretion efficiency of amyloid proteins through the curli biogenesis pathway^38^ (**Fig. 3b**). The engineered *E. coli* strains were cultured in M63 medium with fresh mouse blood at 37 °C adhering strongly to plate surfaces and forming biofilm-like coatings, as shown by Crystal Violet (CV) staining (**Fig. 3c**). Uninduced strains remained suspended with minimal plate adhesion (**Fig. 3c, Supplementary Fig. 5**). Transmission electron microscopy (TEM) images showed extensive self-assembled protein fibers surrounding the bacteria only in the presence of blood (**Fig. 3c**). Congo Red (CR) assays confirmed the amyloid features of these living glue (**Supplementary Fig. 6**). We labeled the living glue comprising CP19K and CP43K as CP19K L-glue and CP43K L-glue, respectively. Sensitivity assays indicated that the living glue system could be activated by blood concentrations as low as 10 ppm (**Fig. 3d**).

To assess adhesion strength, we conducted lap shear tests^39^ (**Fig. 3e**). Strains were cultured on YESCA solid agar plates for two days with 100 ppm blood as an inducer before adhesive force measurements. CP19K and CP43K L-glues displayed strong shear adhesive strength, reaching 330 ± 10 kPa and 404 ± 16 kPa, respectively, surpassing wild-type *E. coli* lacking the adhesive matrix by 6 to 8 times (**Fig. 3e**). Rheological analysis further demonstrated that the storage modulus of both CP19K and CP43K L-glues exceeded that of the wild type control (**Supplementary Fig. 7**), likely due to the cross-linked fibers within the living glue that enable greater mechanical stress resilience.

To evaluate tissue adhesion, engineered and wild-type *E. coli* strains were cultured and induced on porcine intestinal tissue samples (**Fig. 3f**). After a two-day incubation and subsequent washes with sterile water to remove non-adherent bacteria and harsh treatments with EDTA solution, bacterial loads were quantified by CFU counting (**Fig. 3g**). CFU counts for L-glue samples were 10-12 times higher than wild type controls (**Fig. 3g**), consistent with fluorescence data showing substantial tissue attachment in L-glue samples, while wild type *E. coli* exhibited minimal fluorescence (**Supplementary Fig. 8**). These findings indicate that barnacle CP matrices significantly enhance tissue adhesion in engineered microbes.

To evaluate the on-demand adhesion and autonomous sealing capabilities of L-glue, we designed and fabricated a microfluidic device with a 1.2 mm diameter hole in the tube wall to simulate a slightly damaged, bleeding vascular tissue^25^ (**Fig. 3h**). The device was 2 cm in length, 100 mm in width, and 80 mm in height, with inlet and outlet tubing for the continuous flow of the lysed mouse blood. An alumina membrane featuring 200 nm pores was placed over the 1.2 mm hole, allowing the leakage of heme from blood cells into the culture medium (**Fig. 3i**). The device was then immersed in a bacterial culture under a blood flow rate of 10 mL/h. The leakage of heme molecules through the small hole triggered the formation of living glue at the damaged site. Scanning electron microscopy (SEM) revealed that the living glue adhered to and occluded the alumina pores, forming microbial coatings that sealed the damaged area (**Fig. 3i**). In contrast, the wild-type *E. coli* control group showed only sparse microbe residues on the leaking membrane, indicating minimal sealing capacity (**Fig. 3i**). These findings were further supported by Cy3-based leakage experiments, which demonstrated that only devices coated with L-glue were capable of effectively preventing Cy3 fluorescence molecules from escaping the microfluidic channel (**Fig. 3j**). We then measured the burst pressure of the L-glue attached to the microfluidic device. Compared to the non-adhering wild-type *E. coli*, both L-glue samples exhibited substantially higher pressure resistance (**Fig. 3k**). Notably, the burst pressure of CP43K L-glue was approximately three times higher than that of CP19K (**Fig. 3k**), indicating greater mechanical robustness. This enhanced performance may be attributed to its more abundant glue matrix production of CP43K L-glue (**Fig. 3d**).

For *in vivo* testing, wild-type *E. coli* and CP19K/CP43K-secreting strains were administered to gut-bleeding mice. All strains carried miRFP680-expressing plasmids for fluorescence imaging. At 24 hours post-administration, the fluorescence intensity in the colons of CP43K and CP19K L-glue-treated mice was 6-fold higher than in control mice (**Fig. 3l-m**), highlighting the adhesive enhancement of CPs in intestinal tissues. Additionally, we devised a fluorescent probe targeting the 16S rRNA of *E. coli*, employing Fluorescence *in situ* Hybridization (FISH) for the qualitative assessment, localization, and rough quantification of engineered bacteria in *ex vivo* colon tissues. The results confirm increased adhesion of L-glues to the intestinal wall compared to wild-type bacteria (**Fig. 3n-o**), demonstrating the adhesive efficacy of these engineered living glues *in vivo*.

### Hemostatic Effect and Prolonged Treatment with Therapeutic Living Glues

Effective repair of the mucus layer is essential for restoring barrier function and preventing further inflammation in IBD treatment, as mucus integrity underlies gut maintenance^40, 41^. To enhance therapeutic efficacy, TFF3, a mucosal protein and repair peptide from the trefoil factor family, was incorporated into the living glue system. CP43K L-glue, expressing higher glue biomass and adhesion strength than CP19K L-glue (**Fig. 2d–e**), was selected as the basis for these experiments. Hereafter, CP43K L-glue is referred to as L-glue. To create a system that secretes both glue components and TFF3, the CP43K, and TFF3 were sequentially expressed with the CsgA signal peptide. The resulting construct is referred to as therapeutic living glue (TL-glue) (**Fig. 4a**). In parallel, a strain secreting only TFF3 was constructed by replacing the CP genes with TFF3, as illustrated in **Fig. 3b**. This strain, designated as *E. coli* (TFF3), serves as a model for typical engineered microbial therapy. Additionally, a wild-type *E. coli* strain, designated as *E. coli* (WT), was developed as a negative control carrying all the genetic components of TL-glue except for CP43K and TFF3.

**Figure 4:**
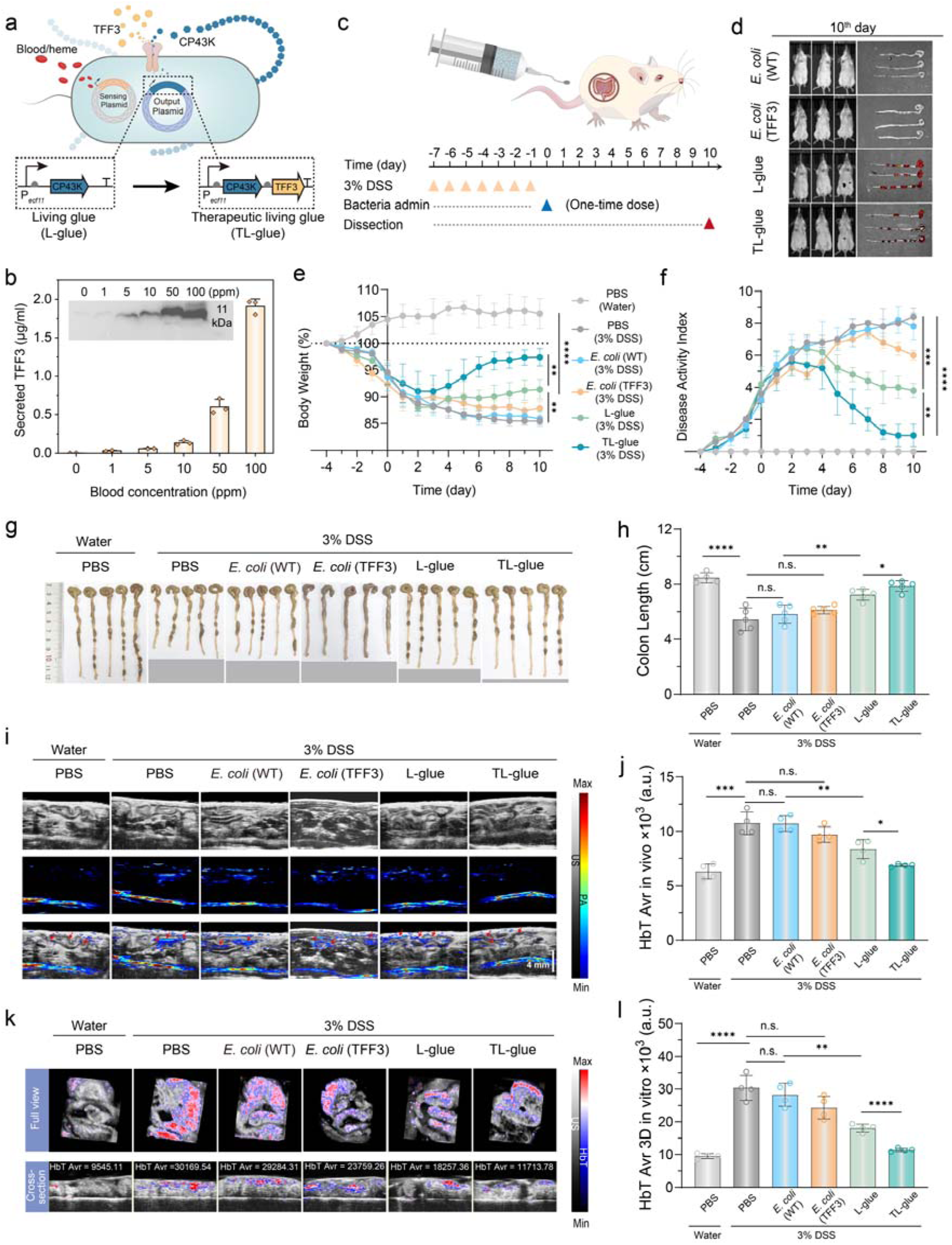
Evaluation of *in vivo* anti-IBD and hemostatic efficacy of living glues. **(a)** Diagram illustrating the genetic circuit design for engineering living glue (L-glue) and therapeutic living glue (TL-glue). **(b)** Dose-response of TFF3 secretion in relation to blood concentration, analyzed via Western blot and quantified by ELISA (*n* = 3 independent experiments, mean ± SD). **(c)** Schematic of the experimental procedure in DSS-induced IBD mice, where 3% DSS was administered 7 consecutive days, followed by a single rectal administration of engineered microbes at a bacterial concentration of OD_600_ = 2. **(d)** Representative *in vivo* fluorescence imaging (FLI) and *ex vivo* colon tissue images from IBD mice 10 days post-administration. **(e)** Body weight changes and **(f)** Disease activity index (DAI) recorded over 10 days post-treatments (*n* = 5 independent experiments, mean ± SD). DAI scoring criteria are listed in **Supplementary Table 1**. **(g)** Digital images (**Supplementary Fig. 12**) and **(h)** colon lengths of *ex vivo* tissues of experimental mice (*n* = 5 independent experiments, mean ± SD). **(i)** Photoacoustic imaging (PAI) and **(j)** semi-quantitative analysis of *in vivo* colon tissues on day 10 post-treatment (*n* = 4 independent experiments, mean ± SD). Top/middle/bottom images display US/PA/Merge modes, respectively. Arrows mark intestinal tissue. Frequency: 40 MHz, Wavelength: 750/850 nm. Scale bar = 4 mm. **(k)** PAI and **(l)** semi-quantitative analysis of *ex vitro* colon tissues 10 days post-treatments (*n* = 4 independent experiments, mean ± SD). n.s. indicates not significant; \**p* < 0.05, \*\**p* < 0.01, \*\*\**p* < 0.001, and \*\*\*\**p* < 0.0001, determined by Student’s *t*-test.

TFF3 secretion efficiency in the engineered TL-glue strain was then evaluated, showing a proportional increase with rising blood concentration (**Fig. 4b**). This dose-response in therapeutic protein production supports precise dosing adjustments to minimize side effects. The long-term colonization of the engineered strains was tested in a mouse model of DSS-induced IBD. Female BALB/c mice received 3% DSS for 7 days to induce colitis (**Fig. 4c**). The L-glue, TL-glue, wild-type *E. coli* control, all expressing similar levels of miRFP680, were compared (**Supplementary Fig. 9**). Over a 10-day fluorescence imaging period, stronger signals were detected in the L-glue and TL-glue groups compared to the *E. coli* (WT) control (**Fig. 4d, Supplementary Fig. 10**). Mice administered with *E. coli* (TFF3), which does not form glue, showed a decline in fluorescence signals over three days, similar to the wild-type *E. coli* control (**Fig. 4d, Supplementary Fig. 10**). The differences in fluorescence signals observed in *ex vivo* gut tissues at the final time point were consistent with the *in vivo* findings (**Supplementary Fig. 11**). These results confirm that glue matrices produced by the engineered bacteria enhance their long-term colonization.

The therapeutic efficacy of living glues was assessed in DSS-induced IBD mice on day 10, where DSS-treated mice showed various IBD symptoms, such as weight loss, an elevated disease activity index (DAI), shortened colons, and tissue damage, confirming the successful establishment of IBD model induction in BALB/c mice (**Fig. 4e-h, Supplementary Fig. 12**). IBD mice were then treated with a single rectal administration of various engineered bacteria. Notable alleviation of DSS-induced IBD was observed in both L-glue and TL-glue groups, as evidenced by weight gain, reduced colon shortening, and decreased colon damage. The TL-glue group showed the most pronounced improvements across all health indices, closely resembling normal group values (**Fig. 4e-h, Supplementary Fig. 12**). This improvement was attributed to the synergistic effects of the CP43K glue’s adhesion properties and the healing capabilities of TFF3. In contrast, the group administrated with *E. coli* (TFF3) only showed mild symptom relief (**Supplementary Fig. 11, 12**), likely due to rapid clearance from the intestine resulting from low tissue adhesion (**Supplementary Fig. 10, 11**).

To examine the hemostatic effects of TL-glue, photoacoustic/ultrasound (PA/US) imaging at 40 MHz and wavelengths of 750/850 nm was used to detect hemoglobin signals. PA signals increased in intestinal tissues of DSS-treated control groups (PBS), confirming successful bleeding induction in the IBD models (**Fig. 4i, j**). Administration of the wild-type *E. coli* strain or the *E. coli* (TFF3) strain yielded nearly no or only limited hemostatic effects, while the L-glue group reduced hemoglobin levels, indicating the notable hemostatic capability of the CP43K-based glue (**Fig. 4i, j**). Weaker PA signals in the TL-glue group (**Fig. 4i, j**) indicated superior hemostatic performance, further verifying the efficacy of TFF3 integration in alleviating IBD symptoms. Variations in hemoglobin content among TL-glue-treated mice were evident in both full-view and cross-sectional images of *ex vivo* colonic tissues (**Fig. 4k, l**), further demonstrating the pronounced hemostatic effects from the combined CP43K glue and TFF3 treatment.

### Histological Analysis and Anti-inflammatory Performance

To further investigate the therapeutic effects of living glues on intestinal inflammation and tissue repair, histological analyses were performed on the colons of treated mice. As shown in **Fig. 5a** and **5b**, mice treated with wild-type *E. coli*, which lacks adhesive, hemostatic, and healing properties, displayed severely distorted crypts, goblet cell depletion, and extensive immune cell infiltration, similar to untreated DSS controls. In contrast, both the *E. coli* (TFF3) and L-glue groups showed improved histopathological conditions with lower histopathological scores compared to the wild-type *E. coli* group, indicating that the adhesion properties of the CP43K glue and the healing capabilities of TFF3 alone can contribute to tissue repair. Notably, the TL-glue group had the lowest histopathological scores, closely resembling those of healthy controls (**Supplementary Table 2)**. Mice in this group showed intact epithelium, preserved crypt architecture, and abundant goblet cells in the distal colon near the anus. These findings highlight the TL-glue’s ability to mitigate intestinal damage and support tissue repair, attributed to prolonged colonization of engineered microbes and continuous therapeutic protein release.

**Figure 5:**
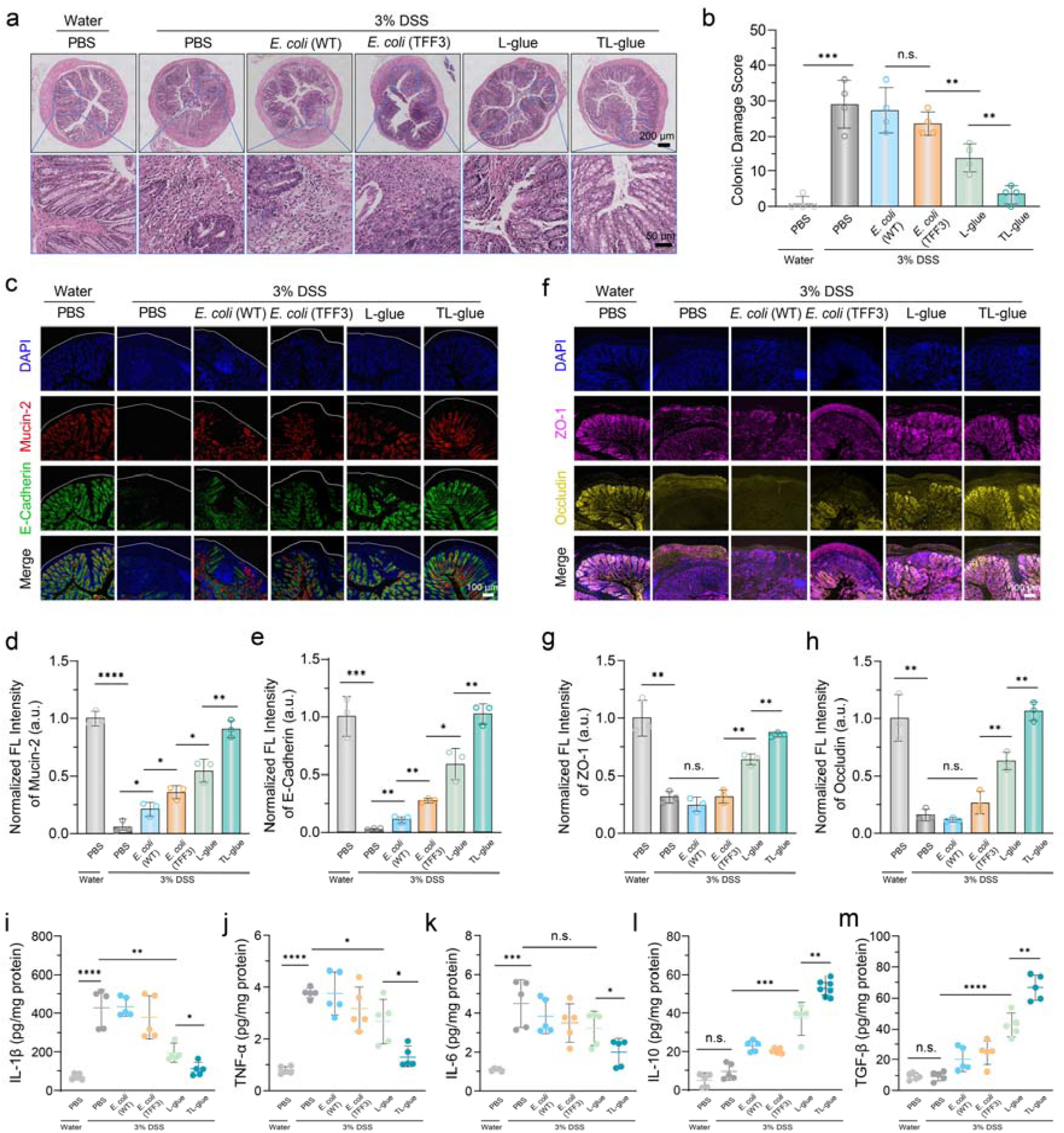
Histological analysis and anti-inflammatory efficacy evaluation of the living glues. **(a)** Representative hematoxylin & eosin (H&E)-stained sections and **(b)** colon damage scores of distal colon tissues from each treatment group (*n* = 4 independent experiments, mean ± SD). The lower panel in **(a)** shows close-up views. Scale bars = 200 and 50 μm, respectively. Scoring criteria for **(b)** are provided in **Supplementary Table 2**. **(c, f)** Representative immunofluorescence staining images and (**d, e, g, h**) semi-quantitative analysis of colon tissue sections post-treatment in mice (*n* = 3 independent experiments, mean ± SD). Blue: DAPI, Red: Mucin-2, Green: E-Cadherin, Pink: ZO-1, Yellow: Occludin, Black: overlay. Scale bar = 50 μm. **(i, j, k, l, m)** Levels of cytokines IL-1β, TNF-α, IL-6, IL-10, and TGF-β from tissue homogenates, determined by ELISA (*n* = 5 independent experiments, mean ± SD). n.s. represents not significant, \**p* < 0.05, \*\**p* < 0.01, \*\*\**p* < 0.001, and \*\*\*\**p* < 0.0001, determined by Student’s *t*-test.

The influence of living glues on the structure and integrity of intestinal epithelial cells was then examined. Specifically, Mucin-2, a key glycoprotein in the intestinal mucus layer, and E-cadherin, essential for cell-cell adhesion epithelial integrity, were analyzed. Immunofluorescence staining revealed slightly elevated levels of Mucin-2 and E-cadherin in colonic tissue treated with *E. coli* (TFF3) and significantly elevated levels in tissues treated with L-glue and TL-glue, with the TL-glue group reaching 90.3% and 100% of the expression levels seen in healthy controls, respectively (**Fig. 5c-e**). Additionally, the levels of ZO-1 and Occludin, two crucial tight junction proteins that help preserve the epithelial barrier^42, 43^, were notably increased in the glue-secreting groups, with the TL-glue group reaching 89.5% and 100% of healthy control values, while almost no significant changes were observed in the *E. coli* (TFF3) group compared to the wild-type *E. coli* group(**Fig. 5f-h**). These results indicate that the synergy of CP43K and TFF3 in TL-glue effectively restored colonic epithelial barrier integrity.

The anti-inflammatory properties of living glues were assessed by measuring pro– and anti-inflammatory markers using enzyme-linked immunosorbent assay (ELISA) in tissue homogenates from treated mice. The untreated colitis group and the groups treated with either *E. coli* (wild) or *E. coli* (TFF3) strains all showed high levels of pro-inflammatory cytokines, including interleukin-1β (IL-1β), tumor necrosis factor-α (TNF-α), and interleukin-6 (IL-6), reflecting severe inflammation in corresponding mice groups (**Fig. 5i-k**). In contrast, treatment with L-glue and TL-glue significantly reduced these pro-inflammatory cytokines while increasing anti-inflammatory markers interleukin-10 (IL-10) and transforming growth factor-β (TGF-β) (**Fig. 5l, m**), underscoring the strong anti-inflammatory effects of the barnacle cement protein-based living glues. These outcomes align with previous *in vitro* findings showing the anti-inflammatory properties of purified barnacle cement proteins^37^ (**Supplementary Fig. 4**).

Finally, the efficacy of TL-glue in IBD mice was confirmed, and a comprehensive biosafety assessment was conducted. Ten days after TL-glue administration in healthy mice, serum and whole blood samples were collected for biochemical analysis. Biochemical markers, including alanine aminotransferase (ALT), aspartate aminotransferase (AST), alkaline phosphatase (ALP), urea nitrogen (UREA), creatinine (CREA), and uric acid (UA), remained within normal ranges, indicating no adverse effects on liver or kidney functions (**Supplementary Fig. 13**). Hematological parameters such as white blood cell count (WBC), lymphocytes (Lymph, Lymph%), monocytes (Mon, Mon%), and neutrophils (Gran, Gran%) counts and percentages, as well as red blood cell count/hematocrit/distribution width (RBC/HCT/RDW), hemoglobin (HGB), mean corpuscular hemoglobin/concentration (MCH/MCHC), and platelet count/mean volume/distribution width (PLT/MPV/PDW) were consistent with those in healthy controls. These results affirm that TL-glue has no adverse effects and exhibits an excellent biosafety profile in mice.

## Discussion

In this study, we designed a gut-resident, bleeding-activated, and autonomously guided adhesive biomaterial that independently addresses ulcerative colitis symptoms. The engineered *E. coli* contains an artificial genetic circuit combining a blood-sensing module with adhesive and therapeutic protein components. This “living” glue, unlike conventional adhesives with static properties, exhibits dynamic capabilities such as bleeding detection, targeted adhesion, and responsive treatment. In a mouse model of IBD, CP43K L-glue, which only secretes adhesive matrices, showed some therapeutic benefit, likely attributed to the anti-inflammatory effects of barnacle cement proteins. However, the living TL-glue, designed to also release therapeutic proteins, provides a notably stronger and longer-lasting effect at only a single dose, likely due to its ability to colonize the gut and deliver sustained protein therapy.

IBD remains a complex disease to treat, prompting ongoing research into therapies, including small molecules, biologics, biomedical materials, genetically engineered probiotics, and microbe-nanomaterial hybrids^44–50^. Our living glue offers unique advantages over existing treatments, particularly by avoiding immune suppression, which can increase the risk of systemic cancer. Instead, our approach directly repairs the mucus layer, providing a safe and efficient healing method. The engineered probiotics actively localize to inflamed sites, delivering targeted treatment with minimal off-target effects. Adhesive features allow therapeutic microbes to stay at the lesion, prolonging contact and interaction with injured tissue. Protein secretion is regulated by disease-specific biomarkers, enabling precise dosing and minimizing side effects. Additionally, this living glue system is adaptable; engineered probiotics can be modified to recognize diverse disease biomarkers, broadening their therapeutic applications. Its self-regulating feature reduces the need for repeated dosing, improving patient compliance and treatment outcomes—a possible explanation for the efficacy of a single dose in the mouse model of IBD.

Through artificially designed gene circuits, we developed a therapeutic living glue with distinctive features for IBD treatment, highlighting the potential of synthetic biology to engineer a new class of programmable biomaterials. This glue integrates disease-sensing, targeted adhesion, and adaptable protein drug release, making it a promising platform for personalized, localized treatments with reduced systemic side effects. This approach holds the potential for addressing complex diseases where precise, sustained treatment is needed. Future studies should focus on testing the living glue in large animal models to validate its targeted adhesion to bleeding sites. Additionally, designing more complex genetic circuits that incorporate multiple disease-specific sensors and logical gates could enable multifaceted, coordinated therapies. While *E. coli* K12 was used in this study due to its non-pathogenic, easily manipulable nature, future versions could employ more widely accepted probiotics, such as *E. coli* Nissle 1917, to increase public acceptance. The adhesive matrix generated by our engineered bacteria could extend the therapeutic duration, and this strategy might be adapted to engineer other probiotic species, such as *Lactobacillus acidophilus*, *Saccharomyces boulardii*, and *Bifidobacterium bifidum,* to form biocompatible extracellular glues.

As biotechnologies and functional materials evolve, integrating synthetic biology with materials science is poised to expand biomaterial innovation. We anticipate that future therapeutic living materials combine highly programmed cellular frameworks with high-performance biological material components, such as anti-inflammatory proteins or anti-aging polysaccharides, to create customizable treatments for specific medical needs.

## Methods

### Plasmid construction

An *E. coli* MG1655 Δ*csg* strain, with all curli-associated genes (csgB, csgA, csgC, csgD, csgE, csgF, csgG) knocked out, was engineered to develop the blood-sensing living glue. Plasmids were assembled using standard molecular biology techniques, specifically Gibson assembly. The blood-sensing gene circuit was divided across two plasmids: the output plasmid and the sensing plasmid. Representative plasmid maps are shown in **Supplementary Figs 14-15**, and detailed plasmid sequences are provided in **Supplementary Table 3 and Table 4**. The heme transporter gene (ChuA) was controlled by the P*_J23107_* promoter, while the heme-sensitive transcriptional repressor gene (HtrR) was under the P*_ProD_* promoter, and the reporter gene (sfGFP) was driven by the P*_ecf11_* promoter on the output plasmid, which also conferred chloramphenicol resistance. The sensing plasmid, containing amplifier modules under the heme-sensitive promoter P*_L_*_(*HrtO*),_ provided ampicillin resistance. In biosensor screening experiments, the promoter for the HrtR protein was modified with various IGEM synthetic promoters using site-directed mutagenesis PCR. Various P*_L_*_(*HrtO*)_ mutants, along with genes for CP19K, CP43K, TFF3, miRFP680, and heme oxygenase, were synthesized by Tsingke Biotechnology Co., Ltd (China).

### Biosensor characterization

After plasmid transformation, colonies were selected from LB (lysogeny broth) agar plates and cultured in LB medium (10 g/L peptone, 5 g/L NaCl, 10 g/L yeast extract) with the appropriate antibiotics at 37 °C. The antibiotics used were chloramphenicol (50 μg/mL) and ampicillin (50 μg/mL). Overnight cultures were diluted 1:100 in fresh LB medium, transferred to a 96-deep well plate (1 mL per well), and incubated at 37 °C with shaking at 1,000 rpm. for three hours. After incubation, heme or blood was added to each well. Cultures were then incubated for another 12 hours, diluted 1:10, and transferred to a 96-well microplate for fluorescence and bacterial concentration measurements using a BioTek microplate reader. The reporter gene sfGFP was measured with excitation and emission wavelengths set of 485 nm and 528 nm, respectively.

### Glue matrix quantification via Crystal Violet staining

Engineered bacteria were incubated overnight at 37 °C with shaking at 220 rpm. Cultures were diluted 1:100 into M63 medium supplemented with 0.2% glucose, 1mM MgSO_, and appropriate antibiotics. To induce glue formation, blood concentrations from 0 to 100 ppm were added to the medium. Cultures were transferred to a 24-well plate for static incubation at 37 °C. For quantification, the M63 medium was removed, and the wells were gently washed with sterile water to remove unattached cells. Each well was then stained with 800 µL of 0.1% aqueous crystal violet solution (Sigma, USA) for 15 minutes at room temperature, and then rinsed repeatedly with ddH_O.Stained biomass attached to the plate was photographed. For quantitative analysis, 800 µL of 30% acetic acid was added to dissolve the dye, followed by a 15-minute incubation at room temperature. A 125 µL aliquot of the solution was transferred to a 96-well microplate, and absorbance was measured at 550 nm using a BioTek microplate reader. A 30% acetic acid solution in water served as the blank control.

### *In vitro* adhesion assay of engineered living glue

Pig intestine tissues, cut into uniform 1 cm x 1 cm pieces, were sterilized with ultraviolet (UV) irradiation. The sterilized tissues were placed at the bottom of the M63 medium containing engineered bacteria. After two days of incubation, the tissues were washed three times with PBS using vortexing to remove unattached bacteria. Fluorescence imaging was performed to visualize bacterial colonization. Tissue samples were treated with 1mM EDTA solution for 30 minutes and vigorously vortexed to dislodge bacteria. The resultant solution was diluted and plated on agar for CFU counting.

### Procedures for fabricating microfluidic devices

The device was fabricated following our previous protocols^25^. Briefly, A silicon mold was fabricated using SU-8 3025 (MicroChem) on a silicon wafer, followed by UV exposure (M365L2-C1, Thorlabs), development, and cleaning. A mixture of PDMS precursor (Dow Corning, USA) and curing agent at a 10:1 ratio was poured onto the mold and cured at 60°C for 4 hours. The cured PDMS slab was then peeled off, ports were created for fluid exchange, and the slab was bonded to a glass slide using oxygen plasma treatment. To seal the middle port, a porous aluminum oxide membrane (200 nm, SHNTI, Mingna Tech) was affixed using half-cured PDMS as an adhesive. The assembled device was baked at 95°C for 30 minutes to ensure stability before use.

### Procedures for on-demand damage repair demonstration using microfluidic devices

The blood-inducible on-demand repair protocol was based on a previously reported method developed in our laboratory^25^. A microfluidic device with a defined leaking site in its channel was immersed in 100 mL of bacterial M63 culture solution within a 140 mm Petri dish. Lysed defibrinated mouse blood (TOPBio) (prepared by diluting blood 1:10 in simulated gastric fluid containing 0.32% pepsin, 0.2% NaCl, and 84 mM HCl, pH 1.2) or a pure hemin solution (prepared by dissolving hemin powder in 1 M NaOH to 25 mM, diluting to 500 µM with ddH_O, and sterilizing with a 0.2 µm filter) was pumped through the device at a flow rate of 10 μL/h. The bacterial culture surrounding the microfluidic device was incubated at 37°C, enabling the bacteria to produce living glue in response to the blood or heme molecules leaking from the channel. To evaluate the repair effects, the M63-bacterial culture solution in the Petri dish was replaced with an equal volume of PBS. A 50 μM Cy3 solution was then pumped through the device at a flow rate of 250 mL/h. Fluorescence leakage was assessed by collecting 200 μL samples every 10 minutes from three fixed sites near the damaged area for quantification of leaked Cy3 molecules. At least three leaky devices were tested under identical conditions, and the repair effects were evaluated consistently following the same protocol.

### Pressure resistance measurements for engineered living glues

Building on our previously reported protocol^25^, the microfluidic device was incubated in 100 mL of M63-bacterial culture solution at 37°C, enabling the bacteria to produce living glue in response to blood or heme molecules leaking from the channel. The pressure resistance of the engineered living glues was measured according to the ASTM F2392-04 standard, a standardized test method for evaluating sealant material burst pressure^51^. Briefly, a syringe was connected to one end of the microfluidic device, while a digital differential pressure gauge (HCJYET, HT-935) was attached to the other end. The sealed channel was then placed into a custom-designed burst pressure apparatus equipped with a pressure gauge for direct measurement.

### Western blot and ELISA assay of therapeutic living glues

To evaluate TFF3 peptide expression at varying blood concentrations, the TL-glue strain was grown in LB medium until OD ≈ 0.6, induced with blood concentrations ranging from 0 to 100 ppm, and incubated at 37 °C with shaking at 220 rpm for one day. Cultures were centrifuged, supernatant filtered through a 0.22-micron membrane and concentrated tenfold using an Amicon Ultra-15 Centrifugal Filter (3 kDa MWCO, Millipore). Concentrated samples underwent gel electrophoresis, followed by transfer to a polyvinylidene fluoride membrane. The membrane was washed with PBST, blocked, and incubated overnights with a 6x-His Tag Mouse Monoclonal Antibody at 4 °C, then with a horseradish peroxidase-conjugated secondary antibody at room temperature for one hour. After further washing, the membrane was analyzed using a luminescence imager.

For protein quantification, 200 µL of the concentrated bacterial culture supernatant was incubated in an ELISA plate at 4 °C overnight, washed, and blocked. It was then incubated with the primary antibody at 37 °C for one hour, washed, and then incubated with the secondary antibody for another hour. After adding 200 µL TMB for 15 minutes at 37° C, the OD_450_ was measured using a BioTek microplate reader and compared against a standard curve of purified TFF3 protein.

### Caco-2 cell culture and biocompatibility assessment

Human intestinal epithelial Caco-2 cells (BNBIO) were cultured in high-glucose Dulbecco’s Modified Eagle Medium (DMEM, Gibco) supplemented with 10% fetal bovine serum (FBS, Gibco) and 1% Penicillin-Streptomycin (Gibco) in an incubator set at 37°C with 5% CO_. For the experiment, 1 × 10^5^ Caco-2 cells were seeded per well in a 24-well culture plate and incubated overnight under the same conditions to form epithelial monolayers. The next day, the monolayers were treated with *E. coli* variants, including wild-type *E. coli* MG1655, L-glue, TL-glue, and *Salmonella enterica*, all suspended in DMEM containing 1% FBS without antibiotics. After 12 hours of incubation, the culture supernatants were collected and analyzed for IL-8 cytokine levels using ELISA kits (BioLegend) following the manufacturer’s protocol.

### RT-qPCR assays for the anti-inflammatory test of purified barnacle cement proteins

Caco-2 cells (BNBIO) were seeded at a density of 10 × 10^5^ cells per well in 6-well plates and cultured for 3 days until reaching approximately 80-90% confluency. The cells were stimulated with 1 µg/mL LPS-SM (DAKEWE), 10 ng/mL IL-1, and 10 ng/mL TNF-α, and were either treated with 100 µg/mL purified barnacle cement protein (CP19K) or left untreated. Incubation was carried out in 6-well plates at 37°C for 24 hours. PBS-treated cells were used as the control group. Total RNA was extracted from Caco-2 cells using the RNAeasy™ Kit (R0027, Beyotime). A total of 1 µg of RNA was reverse-transcribed into cDNA using the HiScript IV All-in-One Ultra RT SuperMix for qPCR (R433-01, Vazyme Biotech). Target genes were amplified using specific primers (**Supplementary Table 5**), with GAPDH serving as the reference gene. The RT-qPCR reaction mixture consisted of 10 µL SsoAdvanced Universal SYBR Green Supermix (Bio-Rad), 2 µL of two-fold diluted cDNA, 0.5 µL of each primer, and sterile distilled water, adjusted to a final volume of 20 µL. Relative gene expression was quantified using the 2^DΔΔCT^ method^52^.

### Animal IBD model and enema administration

Female BALB/c mice (6-8 weeks old) from the Guangdong Provincial Medical Laboratory Animal Center (Guangzhou, China) were housed in individually ventilated cages, with five mice per cage, under controlled conditions (12-hour light/ dark cycle, 23 ± 3 °C, 40-70% humidity). All animal procedures complied with relevant laws and were approved by the Animal Ethics and Welfare Committee (AEWC) of Shenzhen University (AEWC-202300010). To establish the IBD model, mice were given drinking water containing 3% DSS for seven days, followed by a switch to regular water supplemented with ampicillin (50 μg/mL) and chloramphenicol (50 μg/mL). On day 0, mice were fasted and allowed to defecate before being manually restrained in a head-down, tail-up position. A glycerol-lubricated polyethylene tube lubricated was carefully inserted into the anus and held in an elevated position for 2 minutes to prevent drug leakage. Engineered bacteria were cultivated in LB medium (25 g/L) with ampicillin (50 μg/mL) and chloramphenicol (50 μg/mL) at 37 °C until reaching the exponential growth phase (OD_600_ = 0.6). The LB medium was then removed by centrifugation (150 rpm, 5 minutes), and bacteria were resuspended in phosphate-buffered saline (PBS) buffer containing 20% glucose to achieve an OD_600_ of 2.0.

### Evaluation of treatment effects

IBD model mice were divided into 5 groups and rectally administered with PBS, wild-type *E. coli*, CP43K L-glue, TFF3-secreting strain (*E. coli* [TFF3]), or TL-glue (OD = 2.0, 100 μL), while a group of healthy mice received PBS injections (5 mice per group). Body weights, fecal consistency, and presence of blood in stools were recorded daily over a 10-day treatment period to calculate the Disease Activity Index (DAI), with evaluation criteria provided in **Supplementary Table 1**. After the treatment period, the mice were euthanized, and colon tissues were collected for photographic documentation and length measurement. The distal portion of the isolated colon tissue at the end of the treatment was stained with H&E for evaluation of colon damage, with scoring criteria outlined in **Supplementary Table 2**.

### *In vivo* immunofluorescence staining analysis

After 10 days of treatment, murine colon tissues were collected and fixed with 4% paraformaldehyde. Sections of the fixed distal colon tissue were prepared and incubated with primary antibodies against Mucin-2 (Servicebio, Catalog No. GB11344, dilution 1:5000), E-Cadherin (Servicebio, Catalog No. GB12083, dilution 2:1000), ZO-1 (Servicebio, Catalog No. GB111402, dilution 5:1000), and Occludin (Servicebio, Catalog No. GB111401, dilution 3:1000) at 4 °C for 12 hours. After washing away unbound antibodies, sections were incubated with goat anti-different species HRP-conjugated secondary antibodies at room temperature for 2 hours. Cell nuclei were stained with DAPI. Images of colon sections were captured using a scanner (3DHISTECH, model Pannoramic MIDI) and observed using CaseViewer 2.4 software. Semi-quantitative analysis of images was performed using Image J (v1.54).

### Colon tissue inflammatory factor level detection

Preparation and detection followed the standard ELISA protocol. Colon tissues were homogenized (1:10 w/v) with 0.9% saline containing digestive enzymes under ice bath conditions, and the homogenate was centrifuged for 10 minutes at 2,500-3,000 rpm. The supernatant was collected for measurement. The ELISA detection process consisted of coating, blocking, washing, sample addition, incubation, washing, antibody application, incubation, washing, enzyme conjugate addition, incubation, washing, substrate addition, reaction termination, and result measurement. The specific ELISA kits included the Mouse IL-1β ELISA Kit (Servicebio, Catalog No. GEM0002), Mouse TNF-α ELISA Kit (Servicebio, Catalog No. GEM0004), Mouse IL-6 ELISA Kit (Servicebio, Catalog No. GEM0001), Mouse IL-10 ELISA Kit (Servicebio, Catalog No. GEM0003), and Mouse TGF-β1 ELISA Kit (Servicebio, Catalog No. GEM0051).

### Statistical analysis

All experimental results are presented as mean ± SD. Comparisons between two groups were performed using the Student’s *t*-test. Data analysis was conducted using GraphPad Prism 9.5, Excel 2020, ImageJ 1.54, and Living Image 4.2. *p* < 0.05 represented statistically significant (\**p* < 0.05, \*\**p* < 0.01, \*\*\**p* < 0.001, and \*\*\*\**p* < 0.0001; n.s., not significant).

## Supporting information

Supplementary Data.docx

## Acknowledgments

We are grateful to Prof. Timothy K. Lu (Massachusetts Institute of Technology) and Prof. Mark Mimee (University of Chicago) for providing the prototype blood-inducible plasmid Heme-luciferase used in this study. Additionally, we thank Addgene for supplying the plasmid (pXW109 Hg (RS-RinA-E11)2) employed in our research. This work was partially sponsored by the National Key R&D Program of China (2020YFA0908100 to C.Z) and 2020YFA0908800 to J.L.), the National Science Fund for Distinguished Young Scholars (32125023 to C.Z.), the Shenzhen Science and Technology Program (ZDSYS20220606100606013 to C.Z. and KQTD20190929172538530 to P.H.), the National Natural Science Foundation of China (32201105 to B.A. and 32401222 to Q.Z), the Shenzhen Medical Research Fund (A2303072 to Q.Z. and B2302047 to P.H.), the Postdoctoral Fellowship Program of CPSF (GZC20231714 to S.J.), the China Postdoctoral Science Foundation (2024T170582 and 2024M752123 to S.J.). We thank the Instrumental Analysis Center of Shenzhen University (Lihu Campus, China).

## Author contributions

C.Z., B.A. conceived the concept and directed the research. B.A. and C.G. systematically designed and conducted experiments on biosensor optimization and living glue *in vitro* characterization. P.H., S.J., J.L., and X.C. designed and performed all animal experiments. W.Z. fabricated the microfluidic devices with the guidance of Y.L. X.J. kindly performed all the microfluidic experiments. X.D. and Q.Z. carried out partial molecular cloning experiments. Y.X. and P.Y. assisted in characterizing the biocompatibility of recombinant barnacle proteins. B.A., C.Z., J.L., and P.H. wrote the paper with help from all authors.

