## Supplementary Data.docx for "Engineered Living Glues for Autonomous Detection and On-Demand Treatment of Inflammatory Bowel Disease"

**Methods:**

**Congo Red (CR) assay. A** 10 μL aliquot of bacteria solution (OD_600_ = 1.0) was added to YESCA agar plates supplemented with 100 ppm blood solution and appropriate antibiotics, then incubated at 37 °C for 72 hours to form colonies. Colonies were scraped from the agar, mixed with Congo Red solution (25 μg/mL), and incubated for 20 minutes at room temperature. The OD_600_ absorbance was measured to quantify the bacteria, followed by centrifugation, and the OD_480_ of the supernatant was measured. Congo Red depletion rate was calculated by (OD_480(control)_ - OD_480_)/OD_600_, where OD_480(control)_ was the OD_480_ absorbance of CR solution (25 μg/mL)^1^.

**Shear-lap strength measurements.** Living glue samples (0.2 ± 0.02 g) scraped from the YEASCA plate were applied evenly between two stainless steel sheets (10 × 60 × 0.05 mm) with a 1 × 1 cm overlap. The two glued sheets were incubated at 30 °C and 30% relative humidity for 2 hours, then measured using an Instron 5966 testing machine with a 120 N mechanical sensor and two vertical 100 N tensile clamps. The gap distance between two clamps was set at 5 cm for initial loading, and a 5 mm/min tensile speed was used for measurements, except for specific shear speed tests. The ultimate shear adhesive strength was defined as the maximum shear load divided by the overlapped area of glue application.

**Viscoelastic properties assay.** Living glue samples scraped from YESCA plates were applied for rheological testing on a 25 mm diameter cone plate (101 µm gap) using an Anton Paar MCR 302 rheometer. To minimize water evaporation, measurements were conducted in a closed chamber with surrounding pure water. Storage modulus and loss modulus were recorded with a strain-controlled model test^2^, using strain amplitudes from 0.01% to 10% at a constant frequency of 10 rad/s.

***In vivo* blood sensing performance and colonization evaluation.** Mice received rectal administrations of engineered bacteria (OD_600_ = 2.0, 100 μL). FLI and quantitative analysis were performed using the IVIS^®^ Spectrum system (PerkinElmer, USA) before and 2 hours after injection. For colonization assessment, IBD mice were rectally administered with PBS, wild-type *E. coli*, CP43K L-glue, TFF3-secreting strain (*E.coli* [TFF3]), or TL-glue (OD_600_ = 2.0, 100 μL), while healthy mice received only PBS. *In vivo* FLI was monitored at various intervals (0, 2, 8, 12, 24, 48, 72, 120, 168, and 240 hours). At 240 hours post-injection, mice were euthanized, colonic tissues collected, and *ex vivo* FLI performed and quantitative analysis.

***In vivo* adhesion assessment.** All engineered bacteria were tagged with miRFP680 (excitation wavelength: 640 nm, emission wavelength: 680 nm). Rectal injections of *E. coli* or CP43K L-glue (OD_600_ = 2.0, 100 μL) were given to IBD mice. After 24 hours, FLI and quantitative analysis of colon tissues were performed using the IVIS^®^ Spectrum system. Colon tissues were fixed in 4% paraformaldehyde, sectioned, and incubated with a primary antibody against Mucin-2 (Servicebio, catalog number: GB11344, dilution 1:5000) and a fluorescent probe targeting 16S rRNA of *E coli* K-12 MG1655 at 4 °C for 12 hours. Following washing, sections were incubated at room temperature for 2 hours with an HRP-conjugated secondary antibody. Images were obtained using a 3DHISTECH Pannoramic MIDI scanner and observed using CaseViewer 2.4. Semi-quantitative analysis was performed with Image J 1.53t.

***In vivo* safety assessment.** Eight female BALB/c mice (6-8 weeks) were divided into 2 groups (4 mice each) and received rectal administrations of PBS and TL-glue (OD = 2, 100 μL). Blood samples collected on day 10 were sent to Wuhan Servicebio Technology Co., Ltd (China) for routine blood analysis (using Mindray Veterinary Automatic Hematology Analyzer, BC-2800vet) and serum biochemical assessments.

**Supplementary Figures**

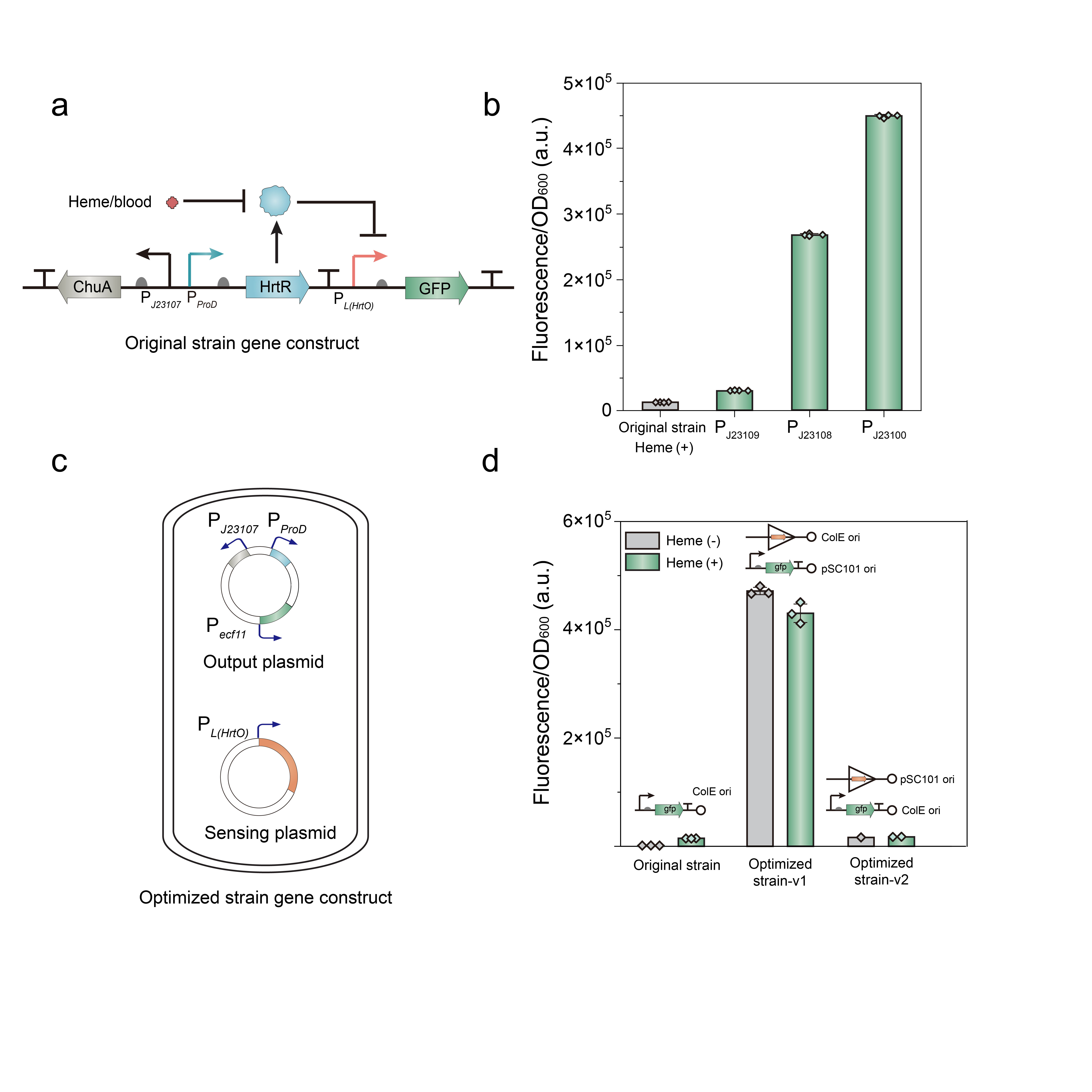

**Supplementary Fig. 1. Optimizing the blood sensor performance using a cascaded transcriptional amplifier circuit.** **a**, Schematic illustrating the genetic circuit design in the original blood-inducible bacteria strain. **b**, The comparison of sfGFP expression of the prototype blood/heme-induced promoter P*_L(HrtO)_* with Anderson promoters P*_J23109_*, P*_J23108_*, and P*_J23100_*. **c**, The cascaded transcriptional amplifier circuit for blood sensing is distributed across two plasmids, the sensing plasmid and the output plasmid. The sensing plasmid contains the amplifier components driven by the blood/heme-responsive promoter P*_L(HrtO)_*. The output plasmid encodes the constitutively expressed proteins ChuA and HrtR, along with an amplifier-specific promoter that regulates the reporter protein sfGFP. **d**, The comparison of the blood-inducible performance of the prototype blood/heme-induced circuit and the early optimized circuits developed in this study. In the first version of the optimized strain (Optimized strain-V1), the amplifier elements are on a high-copy-number ColE1-origin plasmid, and the sfGFP reporter is encoded on a low-copy-number pSC101-origin plasmid. In the second version (Optimized strain-V2), the amplifier elements are on the low-copy-number pSC101-origin plasmid, and the high-copy-number ColE1-origin plasmid is used for the output plasmid. Error bars in the graph indicate the standard deviation from three independent replicates (n = 3).

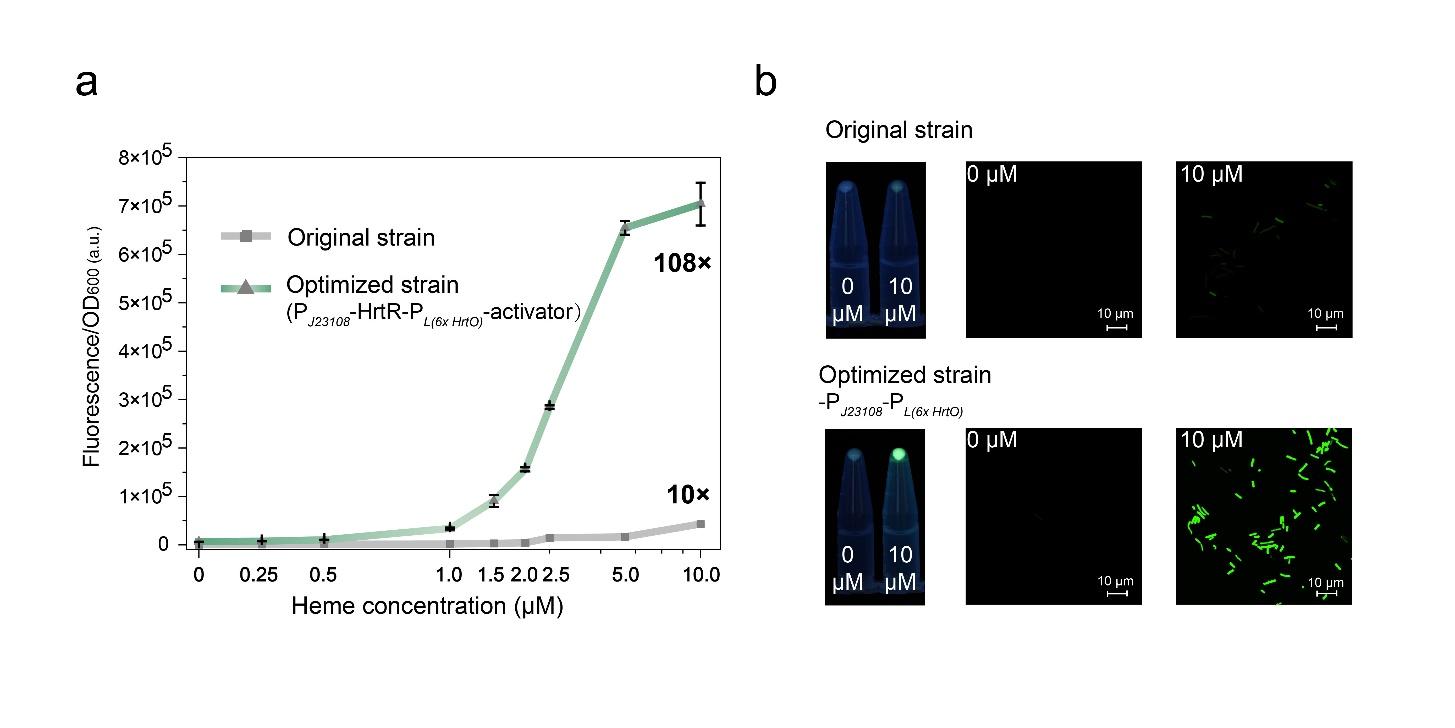

**Supplementary Fig. 2. Comparison of heme-induction performance between the prototype and the final optimized amplifier-integrated circuits. a,** Comparative dose-response analysis of the optimized and prototype strains across varying heme concentrations. Error bars in the graph indicate the standard deviation from three independent replicates (*n* = 3). **b,** Fluorescent images comparing the prototype control and optimized strain.

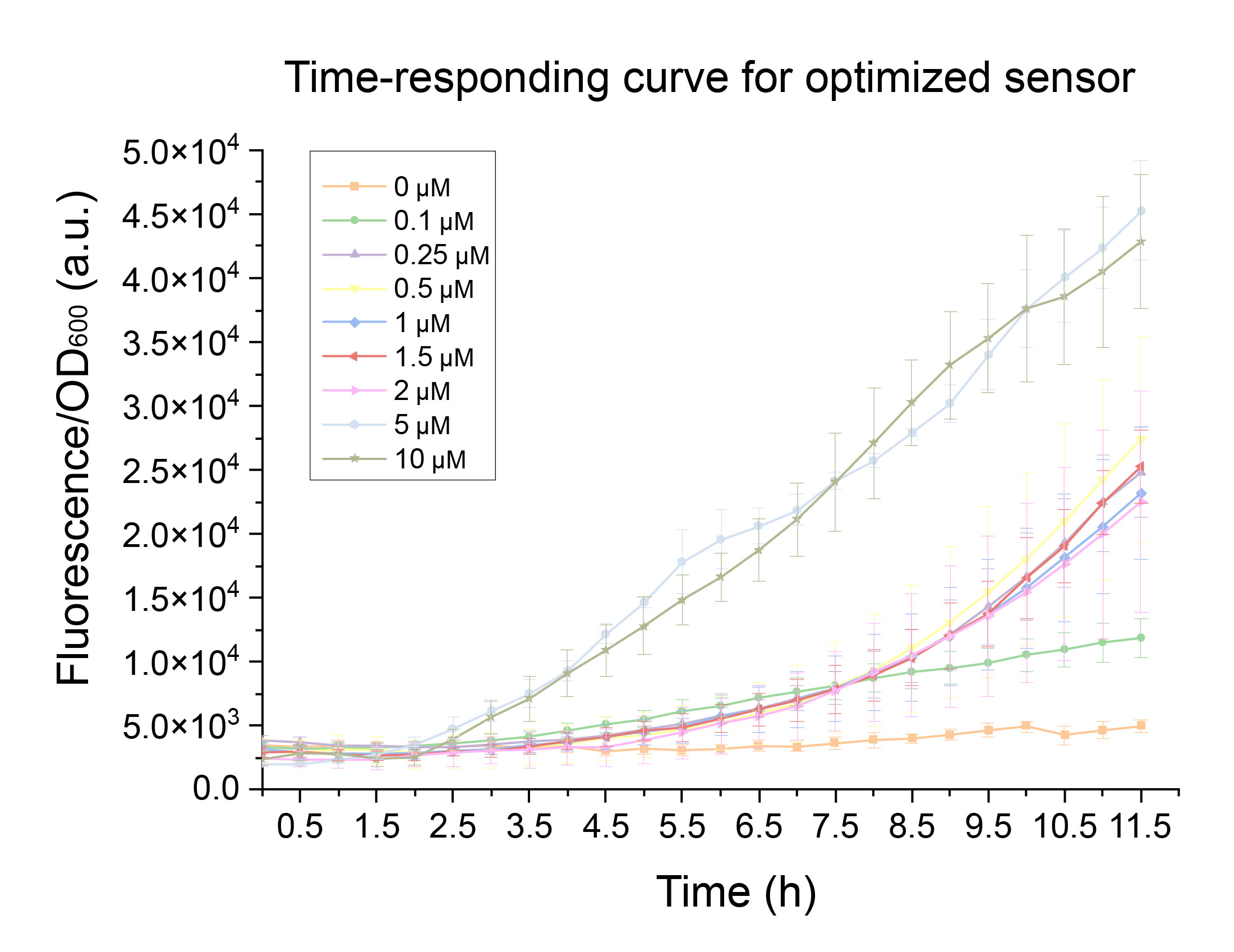

**Supplementary Fig. 3. Time response curve of the optimized blood sensor across various heme concentrations.** The time response curve shows that detectable signals emerge within 2-3 hours post-induction, reaching a peak at 12 hours. Error bars in the graph indicate the standard deviation from three independent replicates (*n* = 3).

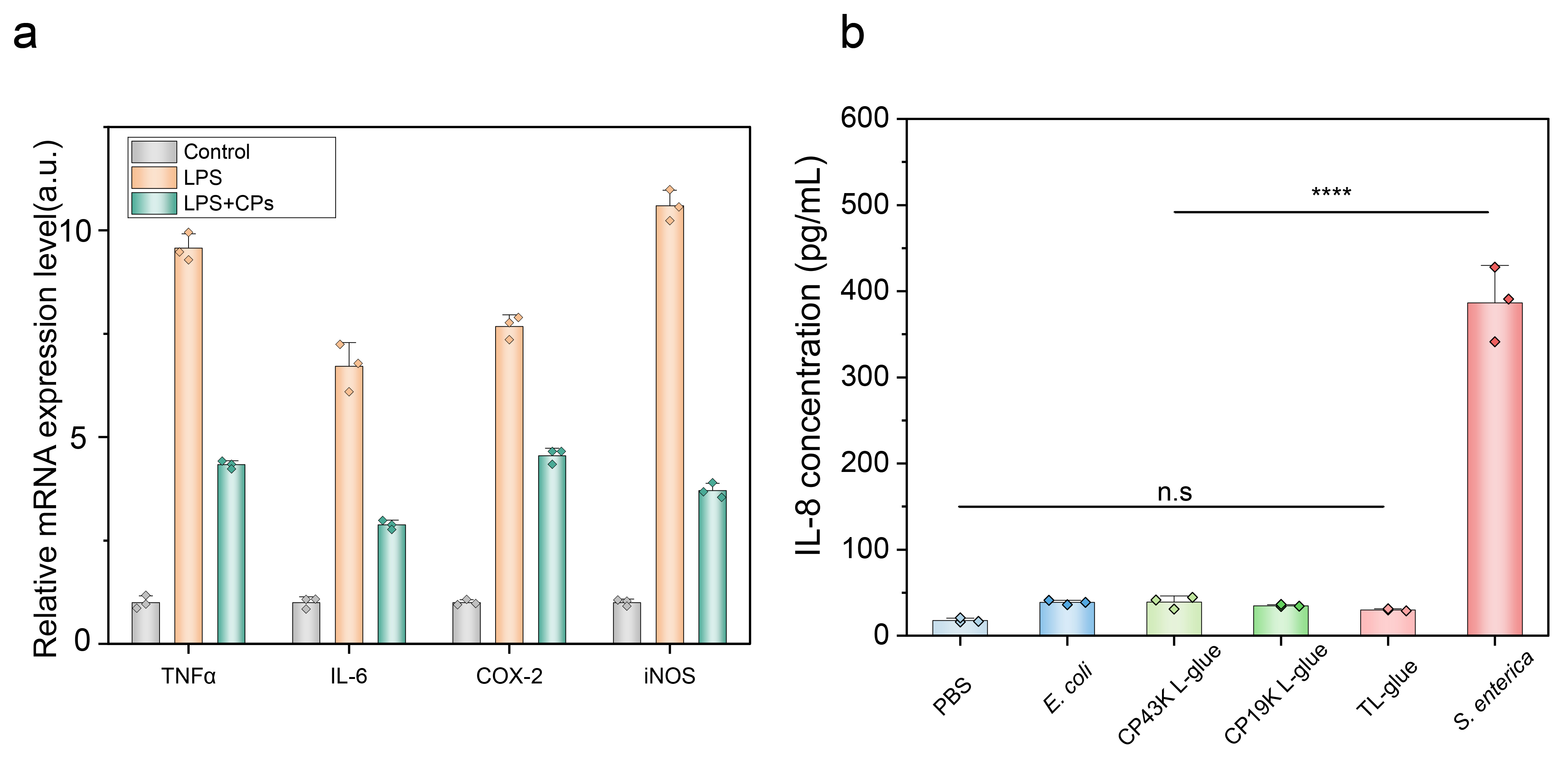

**Supplementary Fig. 4. Anti-inflammatory and biocompatibility tests of the barnacle cement proteins.** **a,** Expression of genes associated with Caco-2 inflammation-related factors measured by RT-qPCR. **b,** Compared to the *Salmonella enterica* treated Caco-2 cells, both the wild-type *E. coli*, CP43K L-glue, CP19K L-glue, and TL-glue samples did not induce significant inflammation, suggesting their biocompatibility and potential safety for human tissue contact.

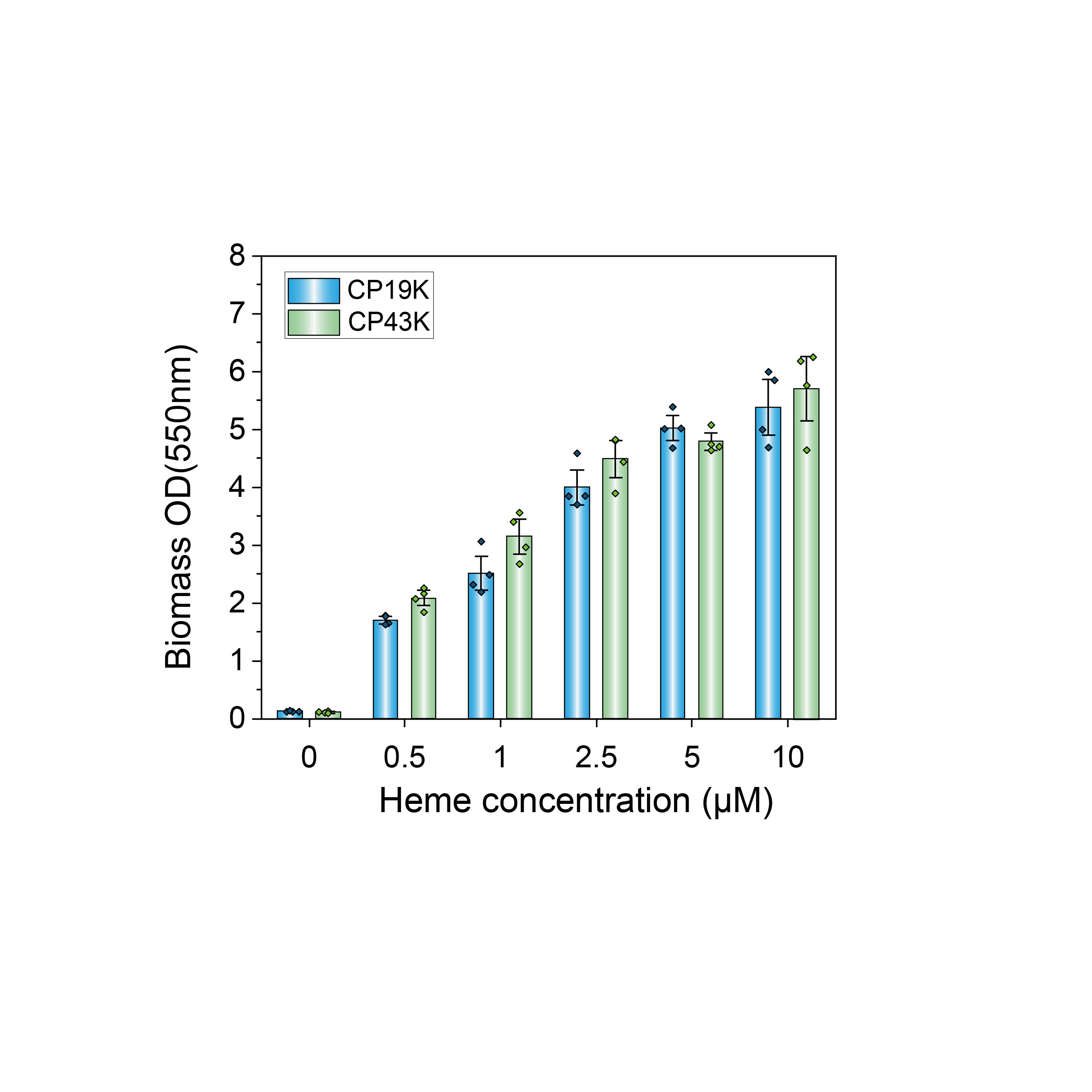

**Supplementary Fig. 5. Quantification of glue biomass in engineered strains via Crystal Violet staining.** Glue biomass production positively correlates with heme concentrations from 0 to 10 μM. Neither the CP19K-secreting strain nor the CP43K-secreting strain formed a glue matrix without heme induction, but both strains produced substantial glue at 0.5 μM. Each column represents data from 4 independent replicates.

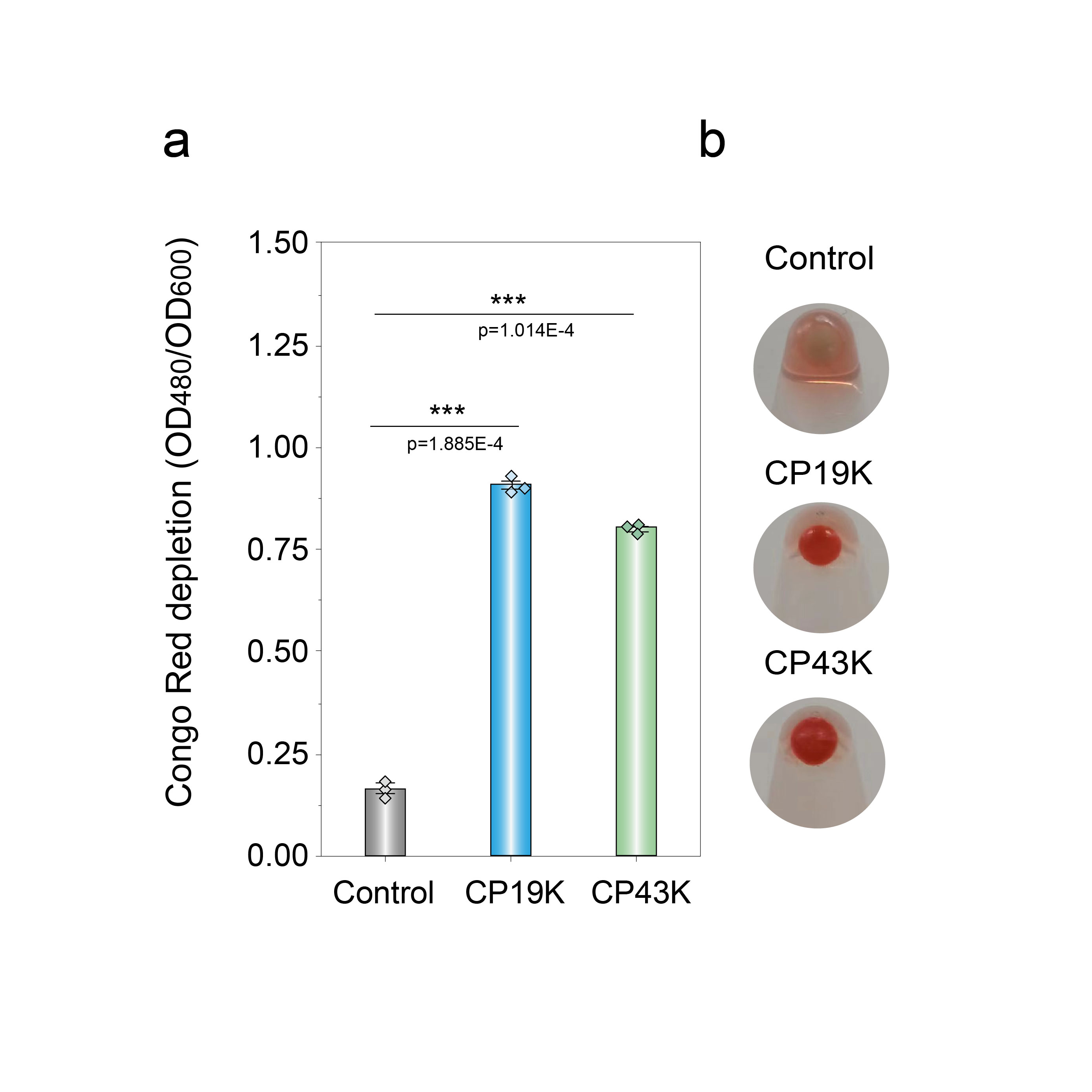

**Supplementary Fig. 6. Quantitative analysis of engineered living glues via Congo Red staining.** **a,** Living glues were prepared from colonies grown on YESCA plates at 37 ℃ for three days with 100 ppm blood induction, confirming amyloid-like features in both CP19K and CP43K recombinant glue matrices through Congo Red staining. Error bars indicate the standard deviation from three independent replicates (*n* = 3). **b,** Representative photographs showing Congo Red-stained engineered bacteria illustrating amyloid characteristics of the living glue materials.

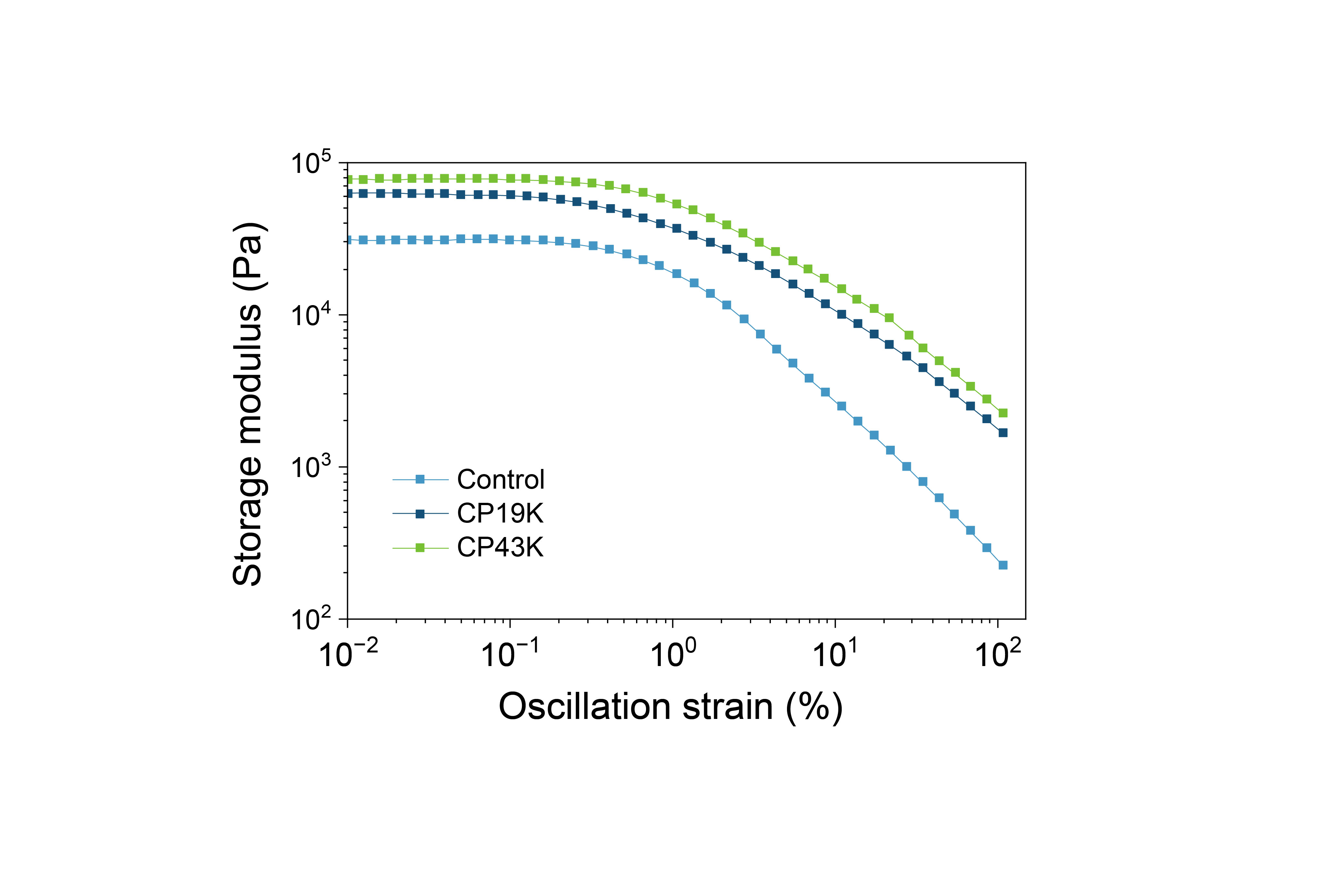

**Supplementary Fig. 7. Storage modulus measurement of engineered living glues via rheology**. Strains were cultured on YESCA plates for three days with 100 ppm blood induction before rheological testing. The storage modulus of engineered living glues, plotted as a function of strain amplitude at a constant frequency (x = 10 rad/s), shows a modest increase due to the proteinaceous extracellular matrices (ECMs).

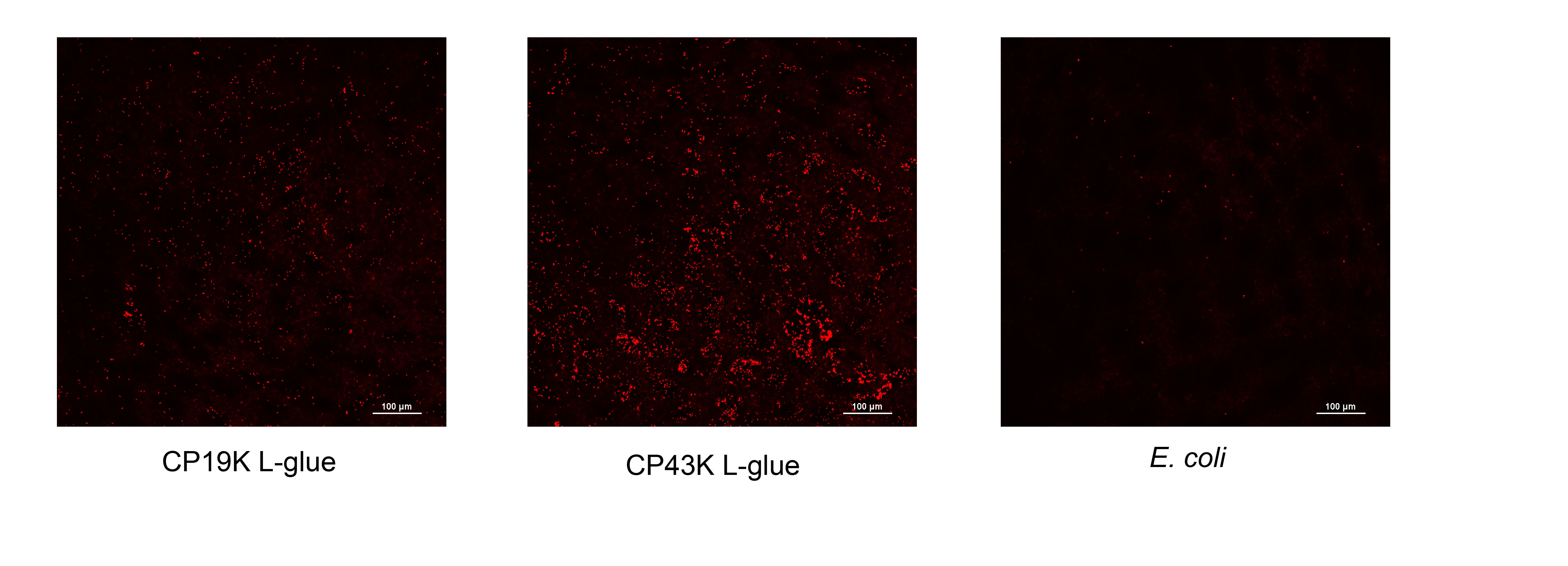

**Supplementary Fig. 8. Fluorescence imaging of strains remaining on gut tissue.** The fluorescence images of gut tissue before detaching the remaining bacteria. The images show substantial tissue attachment in L-glue samples, while wild-type *E. coli* exhibited minimal fluorescence

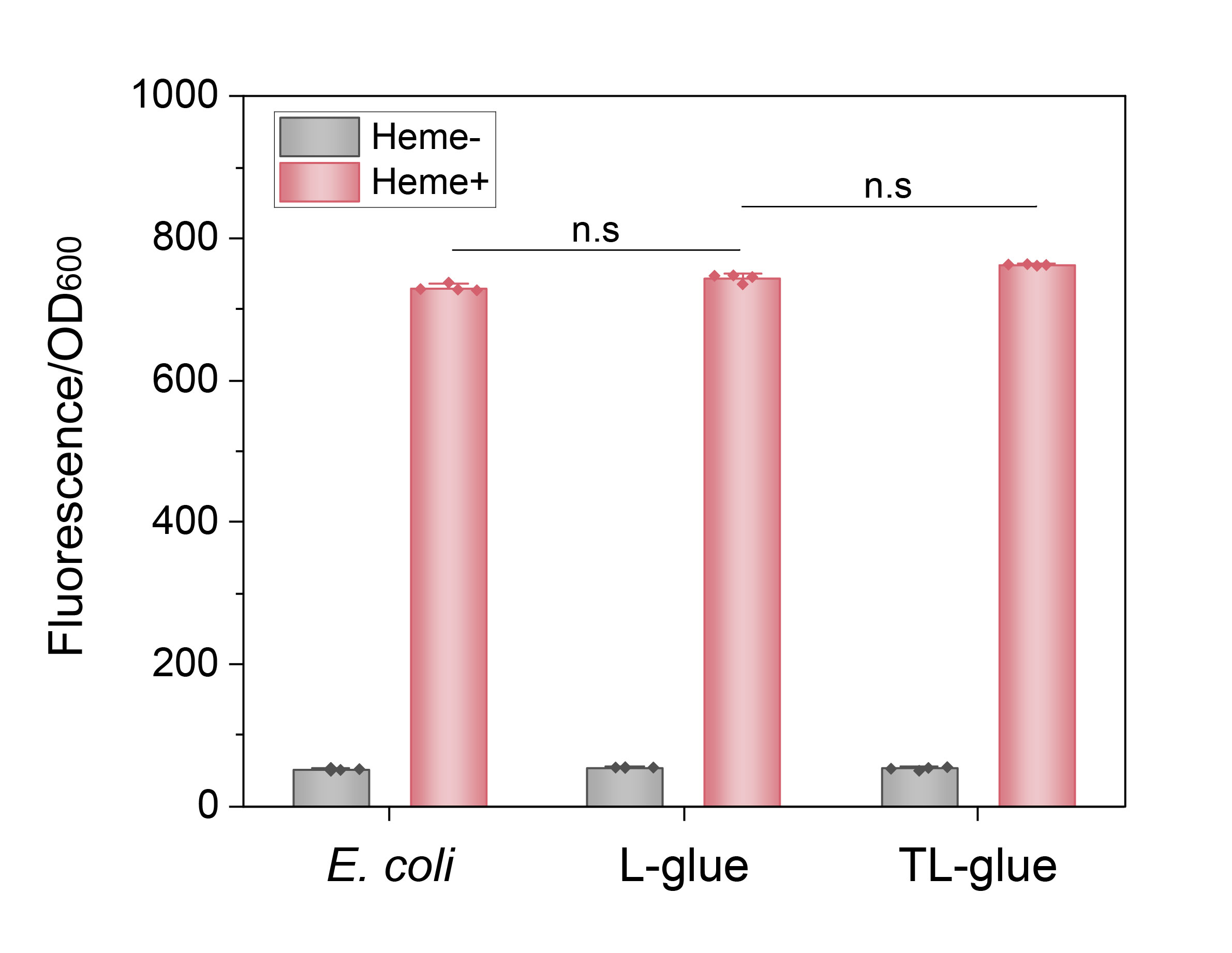

**Supplementary Fig. 9. The L-glue, TL-glue, and engineered *E. coli* control expressing the same level of miRFP680.** To assess fluorescent protein expression levels in engineered *E. coli* strains that encoded a blood-inducible miRFP680-expressing cassette, cultures were grown in LB medium to an OD_600_ of 0.6 and supplemented with heme solution to a final concentration of 5 μM. Fluorescence intensity was measured using a BioTek microplate reader with excitation at 638 nm and emission at 680 nm. The results indicated that the metabolic burden of glue matrix production had minimal impact on miRFP expression.

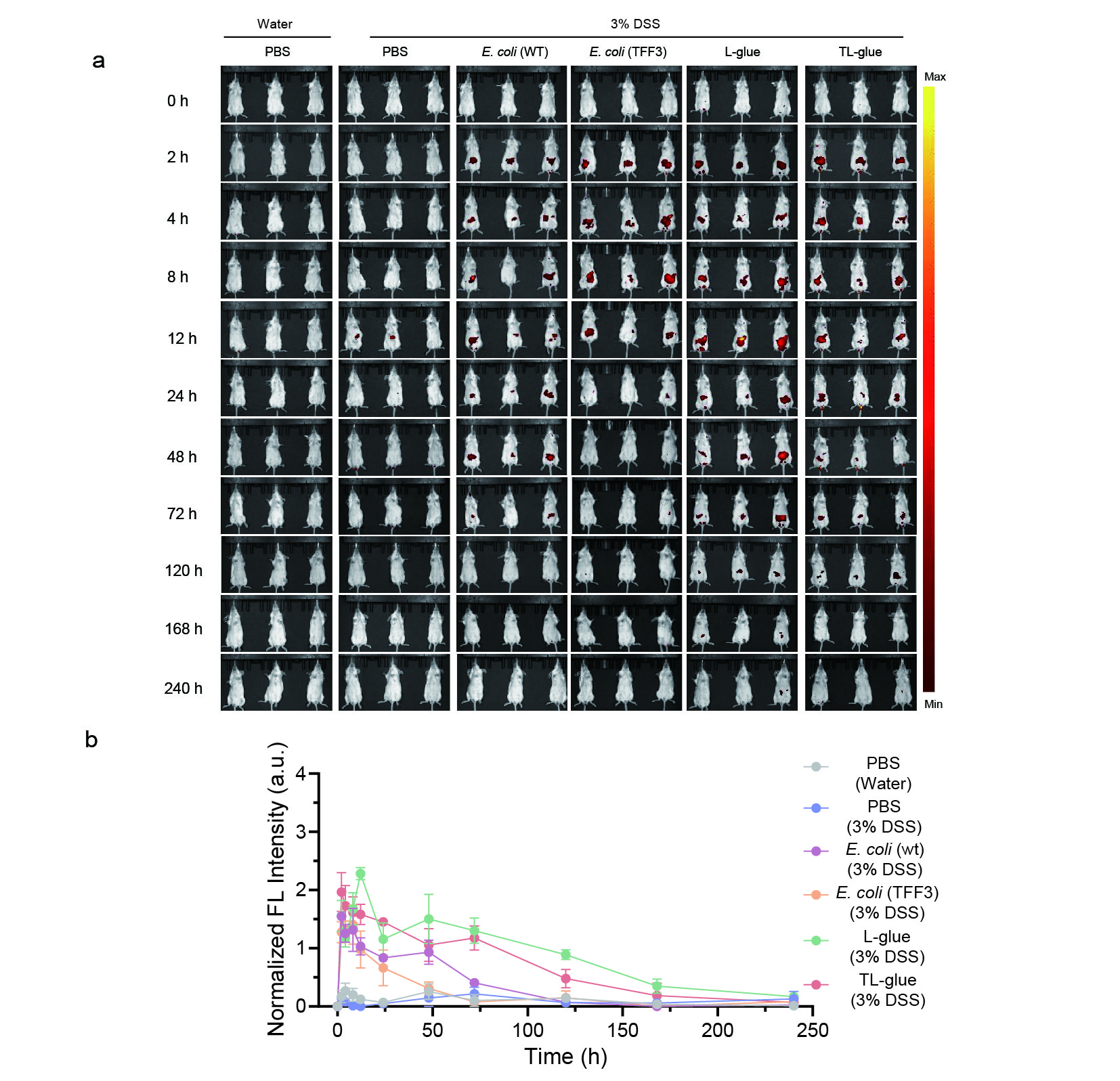

**Supplementary Fig. 10. Colonization efficacy of engineered bacteria in the mouse gut.** **(a)** *In vivo* fluorescence imaging (FLI) images and **(b)** Semi-quantitative analysis over 10 days post-administration of various miRFP680-labeled engineered microbes in mice. Results are presented as mean ± SD (*n* = 3). Statistical significance is represented by n.s. for not significant, **p* < 0.05, ***p* < 0.01, ****p* < 0.001, and *****p* < 0.0001, determined by Student's *t*-test.

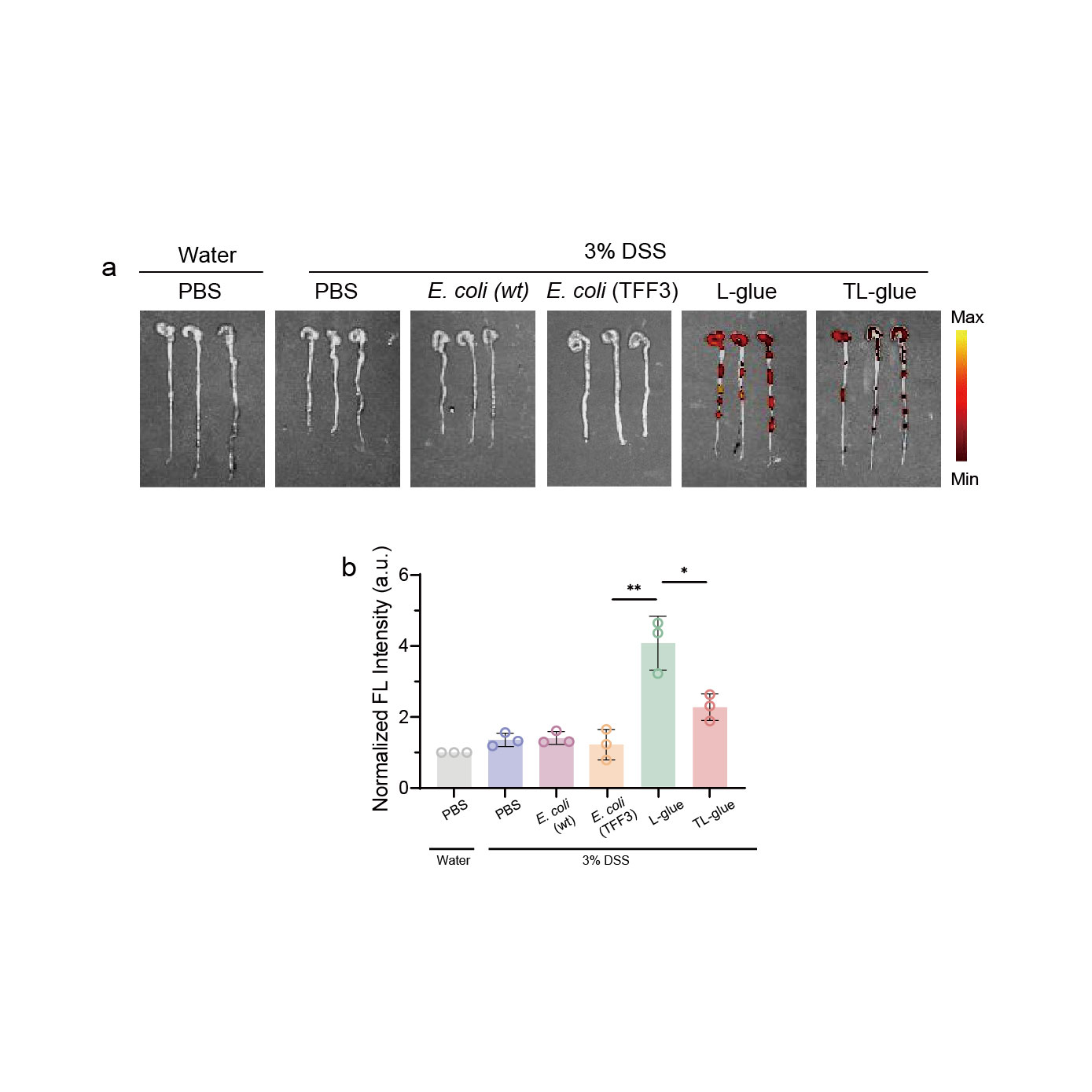

**Supplementary Fig. 11. Colonization efficacy of engineered bacteria in *ex vivo* gut.** FLI images **(a)** and semi-quantitative analysis **(b)** of colonization in *ex vivo* colon tissues on day 10 post-treatment. All results are presented as mean ± SD (*n* = 3 independent experiments). Statistical significance is represented by n.s. for not significant, **p* < 0.05, ***p* < 0.01, ****p* < 0.001, and *****p* < 0.0001, determined by Student's *t*-test.

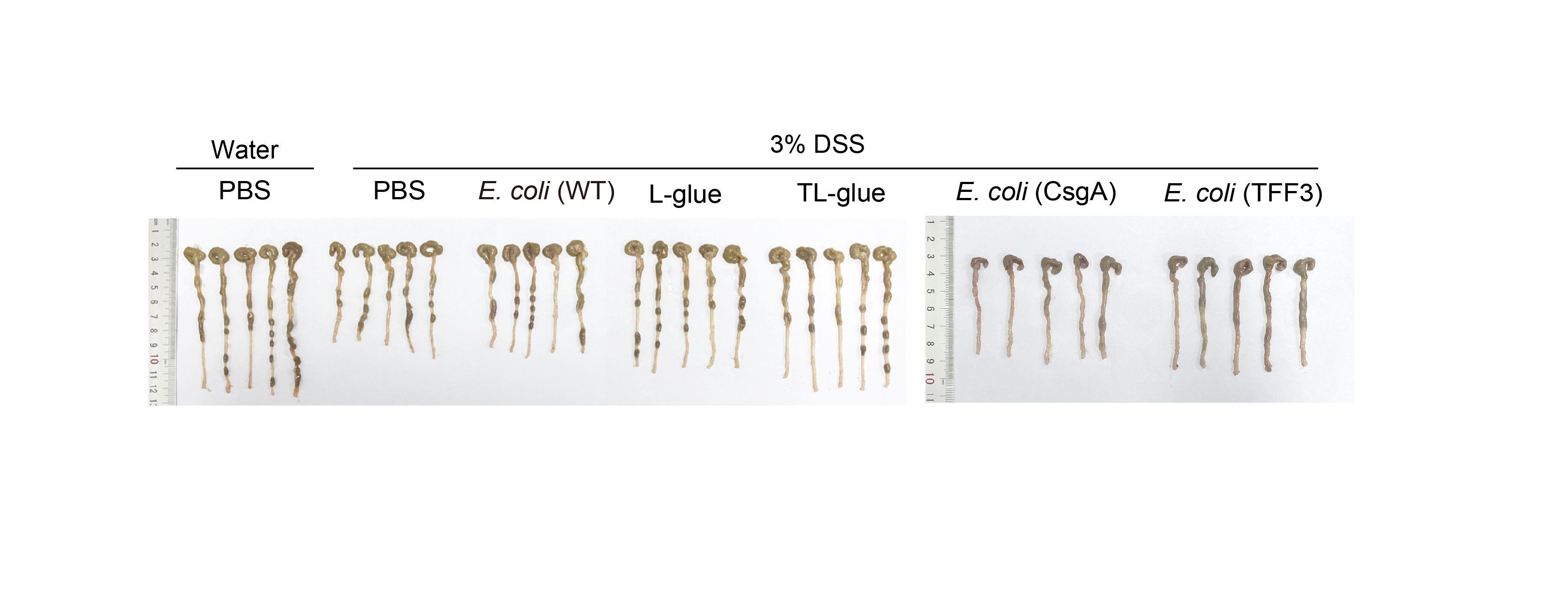

**Supplementary Fig. 12. Colon lengths of *ex vivo* tissues of experimental mice.** The original data showed mouse gut length measurements after 10 days of treatment. An engineered strain that secreted CsgA in response to blood, instead of CP43K or TFF3, was tested for therapeutic performance comparison. The results showed that the CsgA-secreting strain exhibited limited therapeutic effects compared to the L-glue and TL-glue-treated groups.

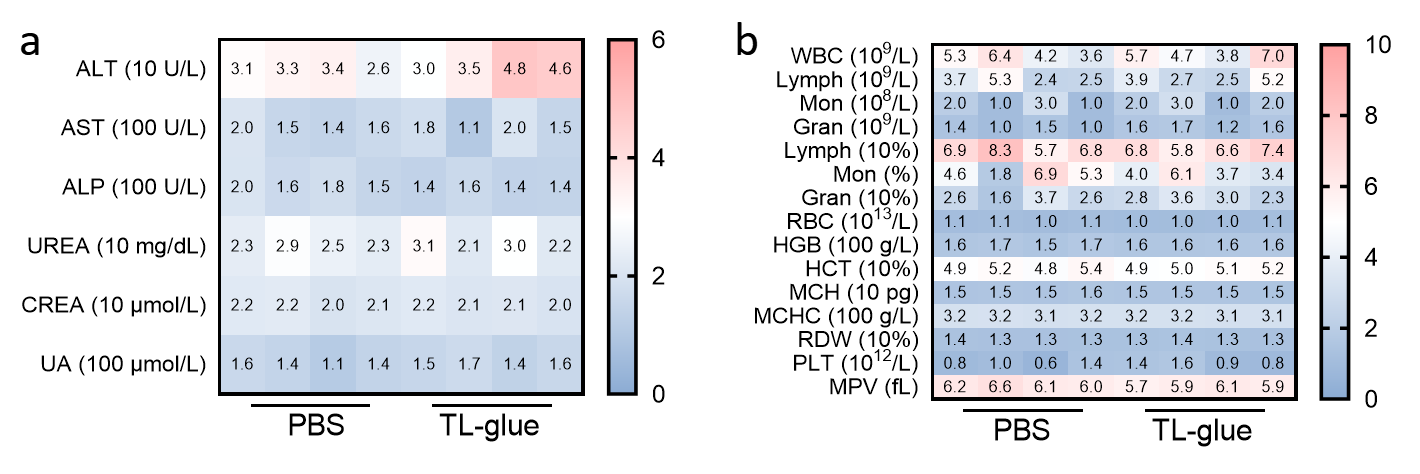

**Supplementary Fig. 13.** **Biosafety evaluation of TL-glue.** Serum **(a)** and hematological biochemical data **(b)** were assessed on day 10 post-TL-glue administration in healthy mice.

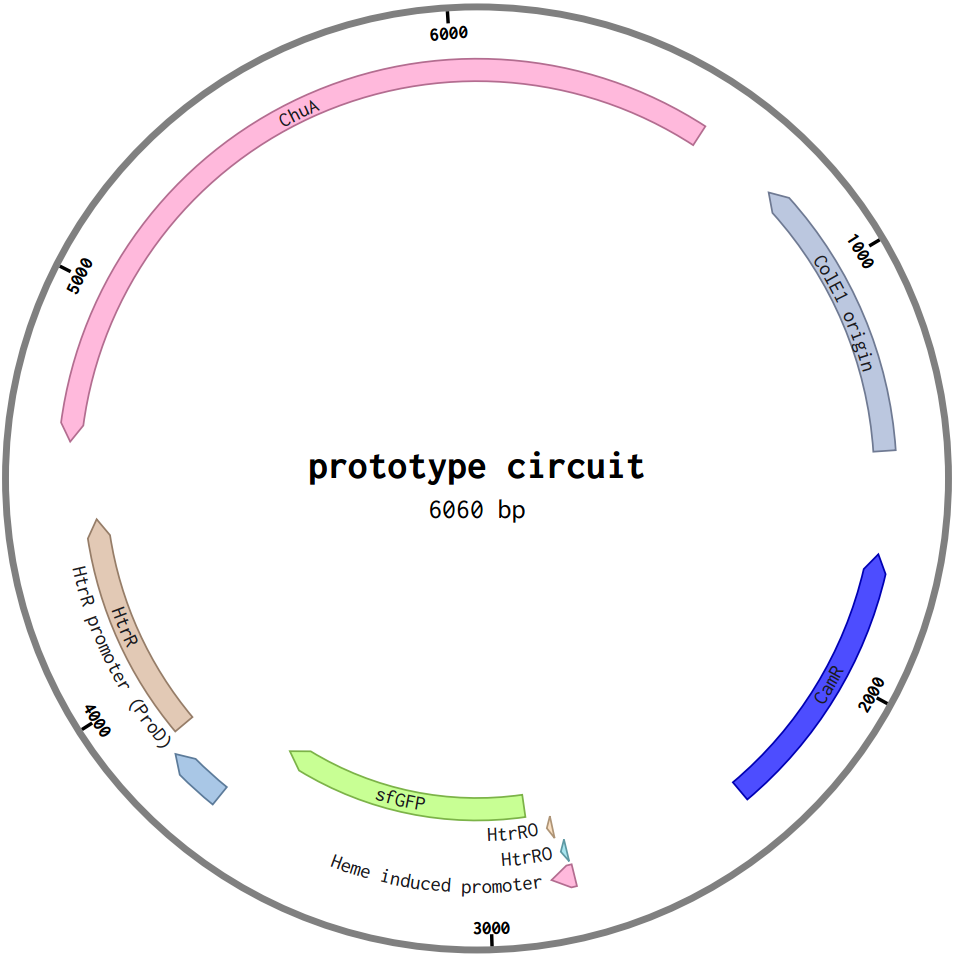

**Supplementary Fig. 14. Plasmid map of the prototyped blood-inducible gene circuit.** Sequence information is provided in Table S2.

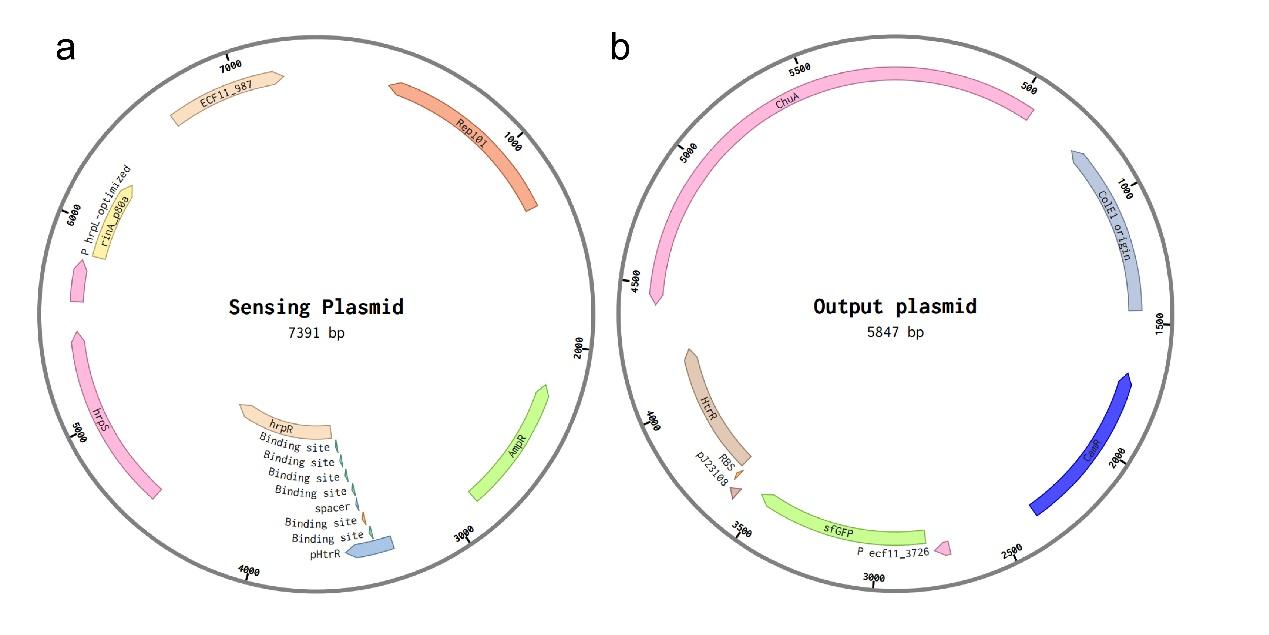

**Supplementary Fig. 15. Plasmid maps of the optimized blood-inducible system.** Maps of the optimized blood-inducible system with both sensing and output plasmids. Detailed sequences are in Table S2.

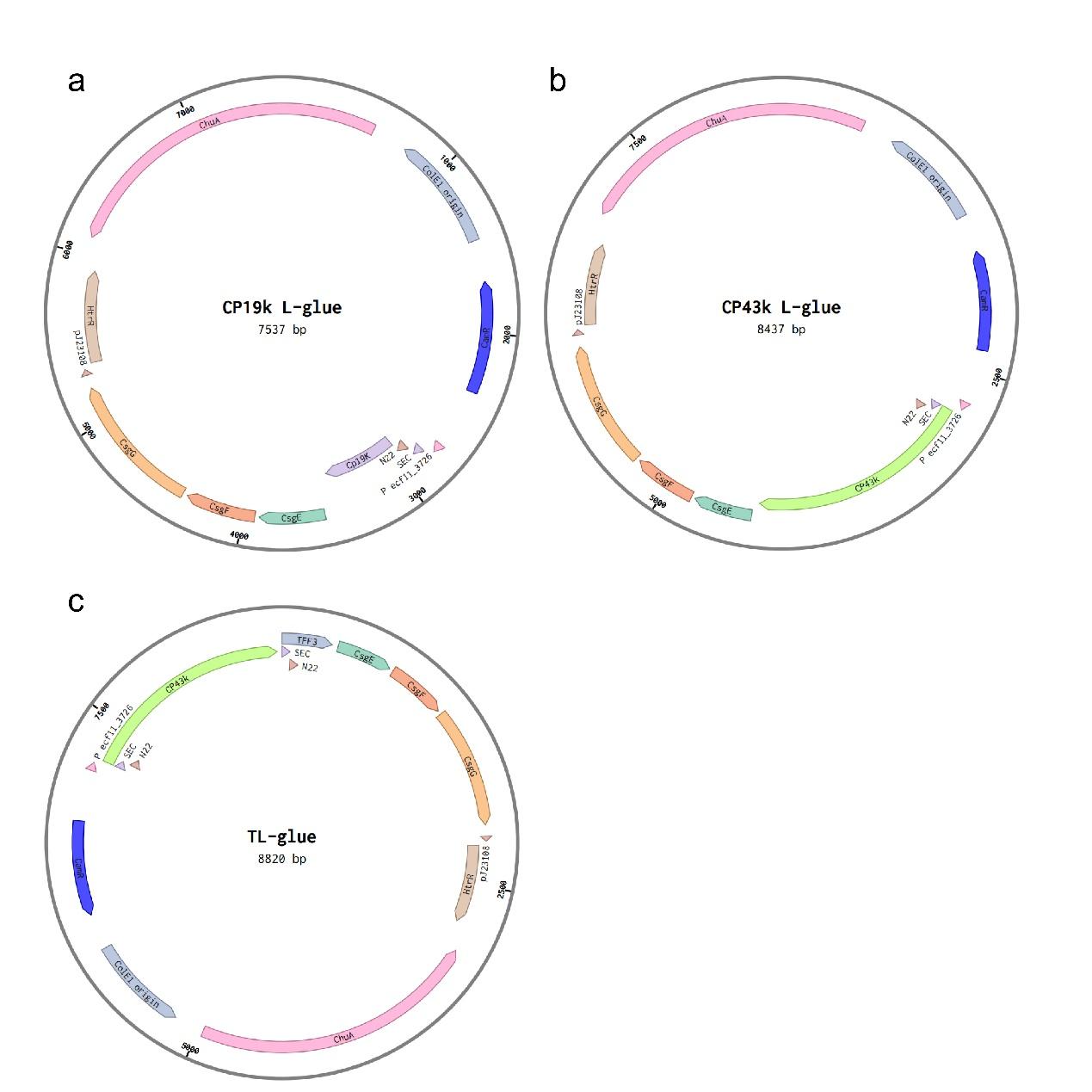

**Supplementary Fig. 16. Plasmid maps of engineered living glues. (a)** Output plasmid map of CP19K L-glue. (**b)** Output plasmid map of CP43K L-glue. **(c)** Output plasmid map of TL-glue. Detailed sequences are in Table S2.

**Table S1: Disease activity index (DAI) parameters and scoring schemes.**

| **Score** | **Weight Loss (%)** | **Stool Consistency** | **Blood in Stool** |
| --- | --- | --- | --- |
| 0 | None | Normal | Normal |
| 1 | 1-5 | Slightly loose stool | Small presence of blood |
| 2 | 5-10 | Loose stool | Significant presence of blood |
| 3 | 10-15 | Diarrhea | Gross blood |
| 4 | > 15 |  |  |

**Table S2: Histological grading scheme for DSS colitis.**

| **Parameter Graded** | **Score** | **Description** |
| --- | --- | --- |
| Severity of inflammation | 0 | None |
|  | 1 | Slight |
|  | 2 | Moderate |
|  | 3 | Severe |
| Depth of injury | 0 | None |
|  | 1 | Mucosa |
|  | 2 | Mucosa and submucosa |
|  | 3 | Transmural |
| Crypt damage | 0 | None |
|  | 1 | Basal one-third damaged |
|  | 2 | Basal two-third damaged |
|  | 3 | Only surface epithelium intact |
|  | 4 | The entire crypt and epithelium lost |
| Percent involvement | ×1 | 0–25% |
|  | ×2 | 26–50% |
|  | ×3 | 51–75% |
|  | ×4 | 76–100% |

**Table S3: DNA sequences of gene elements used in this study.**

| Name | Description | DNA sequence |
| --- | --- | --- |
| P*_J23105_*-RBS | Constitutive promoter | TTTACGGCTAGCTCAGTCCTAGGTACTATGCTAGCTACTAGAGATTAAAGAGGAGAAATACTAG |
| P*_J23108_*-RBS |  | CTGACAGCTAGCTCAGTCCTAGGTATAATGCTAGCTACTAGAGATTAAAGAGGAGAAATACTAG |
| P*_J23107_*-RBS |  | TTTACGGCTAGCTCAGCCCTAGGTATTATGCTAGCACATTTCCAACACTAACCCAAGGGAGCTTTAAATC |
| P*_J23109_*-RBS |  | TTTACAGCTAGCTCAGTCCTAGGGACTGTGCTAGCTACTAGAGATTAAAGAGGAGAAATACTAG |
| P*_J23100_*-RBS |  | CTTGACGGCTAGCTCAGTCCTAGGTACAGTGCTAGCTACTAGAGATTAAAGAGGAGAAATACTAG |
| P*_ProD_*-RBS |  | CACAGCTAACACCACGTCGTCCCTATCTGCTGCCCTAGGTCTATGAGTGGTTGCTGGATAACTTTACGGGCATGCATAAGGCTCGTATAATATATTCAGGGAGACCACAACGGTTTCCCTCTACAAATAATTTTGTTTAACTTTGAATTCTATACTCTAATTAATCACATAATAAGGACGAATTT |
| P_h_*_rpL_*-RBS | Amplifier-specific promoter | GCCGGATTATGTCCGCTGAGTGGGTCACGGTCCCGGATCAGTTCCCTTGCGAAGCTGACCGATGTTTTTGTGCCAAAAGCTGTTGTGGCAAAAAACGGTTTGCGCAAAGTTTTGTATTACAAAGAATTTCACATTTTAAAATATCTTTATAAATCAATCAGTTATTTCTATTTTCAAGCTGGCACGGTTATTGCTATAGGGCTTGTACTACTAGAGATTAAAGAGGAGAAATACTAG |
| P*_rinA_* -RBS |  | AATTGGCAGTAAAGTGGCAGTTTTTGATACCTAAAATGAGATATTATGATAGTGTAGGATATTGACTATCTTACTGCGTTTCCCTTATCGCAATTAGGAATAAAGGATCTATGTGGGTTGGCTGATTATAGCCAATCCTTTTTTAATTTTAAAAAGCGTATAGCGCGAGTACTAGAGTCACACAGGACTACTAG |
| P*_ecf1_*-RBS |  | GCCTCCACACCGCTCGTCACATCCTGTGATCCACTCTTCATCCCGCTACGTAACACCTCTTACTAGAGATTAAAGAGGAGAAATACTAG |
| P*_L_*_(_*_HrtO_*_)_ | The original design of blood inducible promoter | ATAAATGACACAGTGTCATTTGACAAAATGACACAGTGTCATGATACTGAGCACA |
| P*_L_*_(_*_3× HrtO_*_)_ | Optimized design of blood inducible promoter in this study | ATGACACAGTGTCATATAAATGACACAGTGTCATTTGACAAAATGACACAGTGTCATGATACTGAGCACA |
| P *_L_*_(_*_4× HrtO_*_)_ |  | ATGACACAGTGTCATATAAATGACACAGTGTCATTTGACAAAATGACACAGTGTCATGATACTGAGCACATCAGCAGGACGCACTGACCATTATGACACAGTGTCAT |
| P *_L_*_(_*_6× HrtO_*_)_ |  | ATGACACAGTGTCATATAAATGACACAGTGTCATTTGACAAAATGACACAGTGTCATGATACTGAGCACAATGACACAGTGTCATTCAGCATGACACAGTGTCATTGACCATGACACAGTGTCAT |
| HrtR | Transcriptional repressor for the P*_L_*_(_*_HrtO_*_)_ and its optimized variants | ATGCCAAAATCAACCTATTTTAGTCTTTCTGACGAAAAACGAAAACGTGTCTATGATGCCTGTTTACTAGAATTTCAAACGCACTCTTTCCATGAAGCTAAAATCATGCACATCGTAAAAGCACTTGATATCCCAAGAGGAAGTTTTTATCAATACTTTGAAGATTTGAAGGATTCATACTATTATATCTTGTCACAGGAAACTGTCGAGATTCATGATTTATTTTTTAATTTACTAAAAGAATATCCTCTAGAAGTTGCTCTTAATAAATACAAGTATCTTCTTCTTGAAAATTTAGTAAATTCGCCCCAATATAATCTTTATAAATATCGATTTTTAGATTGGACTTATGAATTAGAAAGAGATTGGAAGCCTAAAGGCGAGGTAACTGTTCCCGCTCGTGAACTTGATAATCCTATTTCCCAAGTATTAAAATCAGTCATTCACAATCTAGTTTATCGCATGTTTAGTGAAAATTGGGATGAACAAAAGTTTATTGAAACTTACGATAAAGAAATCAAATTGCTCACAGAGGGCTTGCTTAATTATGTTACTGAAAGCAAAAAATAG |
| ChuA | *E.coli* membrane channel protein that allows the entrance of heme molecules into bacteria | ATGTCACGTCCGCAATTTACCTCGTTGCGTTTGAGTTTATTGGCCTTACCTGTTTCTGCCACCTTGCCAACGTTTGCTTTTGCTACTGAAACCATGACCGTTACGGCAACGGGGAATGCCCGTAGTTCCTTCGAAGCGCCTATGATGGTCAGCGTCATCGACACTTCCGCTCCTGAAAATCAAACGGCTACTTCAGCCACCGATCTGCTGCGTCATGTTCCTGGAATTACTCTGGATGGTACCGGACGAACCAACGGTCAGGATGTAAATATGCGTGGCTATGATCATCGCGGCGTGCTGGTTCTTGTCGATGGTGTTCGTCAGGGAACGGATACCGGACACCTGAATGGCACTTTTCTCGATCCGGCGCTGATCAAGCGTGTTGAGATTGTTCGTGGACCTTCAGCATTACTGTATGGCAGTGGCGCGCTGGGTGGAGTGATCTCCTACGATACGGTCGATGCAAAAGATTTATTGCAGGAAGGACAAAGCAGTGGTTTTCGTGTCTTTGGTACTGGCGGCACGGGGGACCATAGCCTGGGATTAGGCGCGAGCGCGTTTGGGCGAACTGAAAATCTGGATGGTATTGTGGCCTGGTCCAGTCGCGATCGGGGTGATTTACGCCAGAGCAATGGTGAAACCGCGCCGAATGACGAGTCCATTAATAACATGCTGGCGAAAGGGACCTGGCAAATTGATTCAGCCCAGTCTCTGAGCGGTTTAGTGCGTTACTACAACAACGACGCGCGTGAACCAAAAAATCCGCAGACCGTTGGGGCTTCTGAAAGCAGCAACCCGATGGTTGATCGTTCAACAATTCAACGCGATGCGCAGCTTTCTTATAAACTCGCCCCGCAGGGCAACGACTGGTTAAATGCAGATGCAAAAATTTATTGGTCGGAAGTCCGTATTAATGCGCAAAACACGGGGAGTTCCGGCGAGTATCGCGAACAGATAACAAAAGGAGCCAGGCTGGAGAACCGTTCCACTCTCTTTGCCGACAGTTTCGCTTCTCACTTACTGACATATGGCGGTGAGTATTATCGTCAGGAACAACATCCGGGCGGCGCGACGACGGGCTTCCCGCAAGCAAAAATCGATTTTAGCTCCGGCTGGCTACAGGATGAGATCACCTTACGCGATCTGCCGATTACCCTGCTTGGCGGAACCCGCTATGACAGTTATCGCGGTAGCAGTGACGGTTACAAAGACGTTGATGCCGACAAATGGTCATCTCGTGCGGGGATGACTATCAATCCGACTAACTGGCTGATGTTATTTGGCTCATATGCCCAGGCATTCCGCGCCCCGACGATGGGCGAAATGTATAACGATTCTAAGCACTTCTCGATTGGTCGCTTCTATACCAACTATTGGGTGCCAAACCCGAACTTACGTCCGGAAACTAACGAAACTCAGGAGTACGGTTTTGGGCTGCGTTTTGATGACCTGATGTTGTCCAATGATGCTCTGGAATTTAAAGCCAGCTACTTTGATACCAAAGCGAAGGATTACATCTCCACGACCGTCGATTTCGCGGCGGCGACGACTATGTCGTATAACGTCCCGAACGCCAAAATCTGGGGCTGGGATGTGATGACGAAATATACCACTGATCTGTTTAGCCTTGATGTGGCCTATAACCGTACCCGCGGCAAAGACACCGATACCGGCGAATACATCTCCAGCATTAACCCGGATACTGTTACCAGCACTCTGAATATTCCGATCGCTCACAGTGGCTTCTCTGTTGGGTGGGTTGGTACGTTTGCCGATCGCTCAACACATATCAGCAGCAGTTACAGCAAACAACCAGGCTATGGCGTGAATGATTTCTACGTCAGTTATCAAGGACAACAGGCGCTCAAAGGTATGACCACTACTTTGGTGTTGGGTAACGCTTTCGACAAAGAGTACTGGTCGCCGCAAGGCATCCCACAGGATGGTCGTAACGGAAAAATTTTCGTGAGTTATCAATGGTAA |
| HrpR | Transcriptional factor of first layer amplifier | ATGAGTACAGGCATCGATAAGGACGTCCGAGAGTGTTGGGGCGTAACTGCATTATCAGCGGGTCATCAAATTGCAATGAATAGCGCGTTTCTGGATATGGACTTGCTGTTGTGCGGGGAAACCGGCACCGGCAAGGACACACTGGCCAACCGCATTCACGAGTTGTCCAGCAGGTCGGGACCCTTTGTGGGCATGAACTGCGCCGCCATTCCCGAGTCGCTGGCAGAGAGCCAGTTATTCGGTGTGGTCAACGGTGCATTCACCGGCGTATGCCGGGCTCGCGAGGGCTACATAGAGGCCTCCAGTGGTGGCACCTTGTACCTGGATGAAATCGACAGCATGCCGTTGAGCCTGCAAGCCAAACTGCTGCGTGTGTTGGAGAGTCGAGGTATCGAGCGTCTGGGCTCGACCGAATTTATCCCGGTGGATCTGCGGATCATTGCCTCGGCCCAGCGGCCACTGGATGAACTGGTGGAACAAGGACTTTTCCGTCGCGACCTGTTTTTTCGGCTCAACGTGCTGACGCTTCACTTGCCAGCCTTGCGCAAACGTCGTGAACAGATCCTGCCATTGTTCGACCAGTTCACCCAGGGTATCGCTGCCGAGTTCGGACGTCCCGCTCCTGCGCTGGACAGCGGGCGTGTGCAGCTGCTGCTCAGCCACGACTGGCCGGGCAACATCCGCGAATTGAAGTCTGCGGCCAAGCGCTTCGTACTCGGCTTCCCCTTGCTGGGCGCCGACCCTGTGGAAGCGCTTGACCCTGCCACGGGGCTGCGCACGCAAATGCGCATCATCGAGAAAATGCTCATCCAGGATGCCTTGAAGCGGCACAGGCACAATTTCGACGCGGTGCTTCAGGAGTTGGAGTTGCCAAGACGCACCCTGTATCACCGCATGAAGGAACTGGGAGTTGCAGCGCCGATCGCTGCGACGGCCGGGGTCTAA |
| HrpS | Transcriptional factor of first layer amplifier | ATGAGTCTTGATGAAAGGTTTGAGGATGATCTGGACGAGGAGCGGGTTCCGAATCTGGGGATAGTTGCCGAAAGTATTTCGCAACTGGGTATCGACGTGCTGCTATCGGGTGAGACCGGCACGGGCAAAGACACGATTGCCCGACGGATTCATGAGATGTCAGGCCGCAAAGGGCGCCTGGTGGCGATGAATTGCGCGGCCATTCCGGAGTCCCTCGCCGAGAGCGAGTTATTCGGCGTGGTCAGCGGTGCCTACACCGGCGCTGATCGCTCCAGAGTCGGTTATGTCGAAGCGGCGCAGGGCGGCACGCTGTACCTGGATGAGATCGATAGCATGCCGCTGAGCCTGCAAGCCAAATTGCTGAGGGTGCTGGAAACCCGAGCGCTTGAACGGCTGGGTTCGACGTCGACGATCAAGCTGGATATCTGCGTGATCGCCTCCGCCCAATGCTCGCTGGACGACGCCGTCGAGCGGGGGCAGTTTCGTCGCGATCTGTATTTTCGCCTGAACGTCCTGACACTCAAGCTTCCTCCGCTACGTAACCAGTCTGATCGCATAGTTCCCCTGTTCACACGTTTTACGGCCGCCGCCGCGAGGGAGCTCGGTGTTCCCGTTCCCGATGTTTGCCCACTGCTGCACAAAGTGCTGCTGGGCCACGACTGGCCCGGCAATATCCGTGAGCTCAAGGCGGCAGCCAAACGCCATGTGCTGGGTTTCCCCTTGCTGGGCGCCGAGCCGCAGGGCGAAGAGCACTTGGCCTGTGGGCTCAAATCGCAATTGCGAGTGATCGAAAAAGCCCTGATTCAGGAGTCGCTCAAGCGCCACGACAATTGTGTGGATTCGGTAAGCCTGGAACTGGACGTGCCACGCCGTACGCTCTATCGACGCATCAAAGAATTGCAGATCTAA |
| RinA | Transcriptional factor of second layer amplifier | ATGACTAAAAAGAAATATGGATTAAAATTATCAACAGTTCGAAAGTTAGAAGATGAGTTGTGTGATTATCCTAATTATCATAAGCAACTCGAAGATTTAAGAAGTGAAATAATGACACCATGGATTCCAACAGATACAAATATAGGCGGGGAGTTTGTACCGTCTAATACATCGAAAACAGAAATGGCAGTAACTAATTATCTTTGTAGTATACGAAGAGGTAAAATCCTTGAGTTTAAGAGCGCTATTGAACGTATAATCAACACATCAAGTAGGAAAGAACGCGAATTTATTCAAGAGTATTATTTTAATAAAAAGGAATTAGTGAAAGTTTGTGATGACATACACATTTCTGATAGAACTGCTCATAGAATCAAAAGGAAAATCATATCCAGATTGGCGGAAGAGTTAGGGGAAGAGAGGCCTGCTGCAAACGACGAAAACTACGCTGCATCAGTTTAA |
| ECF11 | Transcriptional factor of third layer amplifier | ATGATGAGCGATAGTCCGCAGAAACTGGGTCGTAATGAATGGAATGCATATATGGATAAAGTGAAAGCCAAAGATCGTGAAGCCTTTGCCTTTGTGTTTCGTTTTTATGCACCGAAACTGAAACAGTTCGCCTATAAACATGTTGGCAATGAACAGGTTGCAATGGAAATGGTTCAAGAAACCATGGCAACCGTTTGGCAGAAAGCACATCTGTATGATGGTAAAAAAAGCGCACTGAGCACCTGGATTTATACCATTATTCGTAACCTGTGCTTTGACCTGCTGCGTAAACAGAAAGGTAAAGAACTGCATATTCACAGCGACGATATTTGGCCGAGCGAATATTATCCGCCTGATATGGTTGATCATTATAGTCCGGAACAGGATATGCTGAAAGAACAGGTGGTTAAATTTCTGGATATCCTGCCGAAAAATCAGCGTGATGTTCTGCAAGCAGTTTATCTGGAAGAACTGCCGCATCAGCAGGTTGCAGAACTGTTTGATATTCCGCTGGGCACCGTTAAAAGCCGTCTGCGTCTGGCAGTTGAAAAACTGCGTCATAGCATGCATACCGAACAGCTGTAA |
| sfGFP | Green fluorescent protein | ATGCGTAAAGGCGAAGAGCTGTTCACTGGTGTCGTCCCTATTCTGGTGGAACTGGATGGTGATGTCAACGGTCATAAGTTTTCCGTGCGTGGCGAGGGTGAAGGTGACGCAACTAATGGTAAACTGACGCTGAAGTTCATCTGTACTACTGGTAAACTGCCGGTTCCTTGGCCGACTCTGGTAACGACGCTGACTTATGGTGTTCAGTGCTTTGCTCGTTATCCGGACCATATGAAGCAGCATGACTTCTTCAAGTCCGCCATGCCGGAAGGCTATGTGCAGGAACGCACGATTTCCTTTAAGGATGACGGCACGTACAAAACGCGTGCGGAAGTGAAATTTGAAGGCGATACCCTGGTAAACCGCATTGAGCTGAAAGGCATTGACTTTAAAGAGGACGGCAATATCCTGGGCCATAAGCTGGAATACAATTTTAACAGCCACAATGTTTACATCACCGCCGATAAACAAAAAAATGGCATTAAAGCGAATTTTAAAATTCGCCACAACGTGGAGGATGGCAGCGTGCAGCTGGCTGATCACTACCAGCAAAACACTCCAATCGGTGATGGTCCTGTTCTGCTGCCAGACAATCACTATCTGAGCACGCAAAGCGTTCTGTCTAAAGATCCGAACGAGAAACGCGATCATATGGTTCTGCTGGAGTTCGTAACCGCAGCGGGCATCACGCATGGTATGGATGAACTGTACAAATG |
| miRFP680 | Far-red light excited fluorescent protein^3^ | ATGGCGGAGGGCAGCGTGGCGCGTCAGCCGGATCTGCTGACCTGTGATGATGAACCGATCCACATTCCGGGCGCGATCCAGCCGCACGGTCTGCTGCTGGCGCTGGCAGCGGATATGACCATCGTGGCGGGCTCTGATAACCTGCCGGAACTGACCGGCCTGGCAATCGGTGCGCTGATCGGTCGTTCCGCGGCGGATGTTTTCGATTCTGAAACCCATAACCGTCTGACCATCGCGCTGGCGGAACCGGGCGCGGCGGTTGGTGCGCCGATCACCGTTGGCTTCACCATGCGTAAAGACGCGGGCTTTATCGGTAGCTGGCACCGTCACGATCAGCTGATCTTTCTGGAACTGGAACCGCCGCAGCGCGATGTGGCGGAACCGCAAGCGTTCTTCCGTCGTACCAACTCTGCGATTCGTCGTCTGCAAGCGGCGGAAACCCTGGAAAGCGCGTGCGCGGCTGCGGCGCAGGAAGTTCGTAAAATCACCGGCTTCGATCGTGTTATGATCTATCGTTTCGCGTCTGACTTCTCCGGCGAAGTGATCGCGGAAGATCGTTGCGCAGAAGTTGAATCTAAACTGGGTCTGCACTATCCGGCGTCCACCGTCCCGGCCCAGGCGCGCCGTCTGTATACCATCAACCCGGTTCGTATTATCCCGGATATTAACTATCGTCCGGTTCCAGTAACCCCGGATTTAAACCCGGTGACCGGTCGTCCGATTGATCTGAGCTTTGCTATCCTGCGTTCTGTTAGCCCGTGCCACCTGGAATTTATGCGTAACATCGGTATGCATGGTACCATGAGCATCTCTATCCTGCGTGGTGAACGTCTGTGGGGTCTGATCGTTTGCCACCACCGTACTCCGTACTATGTTGACCTGGACGGGCGCCAGGCATGCAAACGTGTAGCTGAGCGTCTGGCCACCCAAATCGGTGTTATGGAGGAATAA |
| Heme-oxygenase | Enzyme that converts heme into biliverdin | ATGAGTGTCAACTTAGCTTCCCAGTTGCGGGAAGGGACGAAAAAATCCCACTCCATGGCGGAGAACGTCGGCTTTGTCAAATGCTTCCTCAAGGGCGTTGTCGAGAAAAATTCCTACCGTAAGCTGGTTGGCAATCTCTACTTTGTCTACAGTGCCATGGAAGAGGAAATGGCAAAATTTAAGGACCATCCCATCCTCAGCCACATTTACTTCCCCGAACTCAACCGCAAACAAAGCCTAGAGCAAGACCTGCAATTCTATTACGGCTCCAACTGGCGGCAAGAAGTGAAAATTTCTGCCGCTGGCCAAGCCTATGTGGACCGAGTCCGGCAAGTGGCCGCTACGGCCCCTGAATTGTTGGTGGCCCATTCCTACACCCGTTACCTGGGGGATCTTTCCGGCGGTCAAATTCTCAAGAAAATTGCCCAAAATGCCATGAATCTCCACGATGGTGGCACAGCTTTCTATGAATTTGCCGACATTGATGACGAAAAGGCTTTTAAAAATACCTACCGTCAAGCTATGAATGATCTGCCCATTGACCAAGCCACCGCCGAACGGATTGTGGATGAAGCCAATGACGCCTTTGCCATGAACATGAAAATGTTCAACGAACTTGAAGGCAACCTGATCAAGGCGATCGGCATTATGGTGTTCAACAGCCTCACCCGTCGCCGCAGTCAAGGCAGCACCGAAGTTGGCCTCGCCACCTCCGAAGGCTAA |
| CsgE | Curli secretion-associated proteins | ATGAAACGTTATTTACGCTGGATTGTGGCGGCAGAATTTCTGTTCGCCGCAGGGAATCTTCACGCCGTTGAGGTAGAAGTCCCGGGATTGCTAACTGACCATACTGTTTCATCTATTGGCCATGATTTTTACCGAGCCTTTAGTGATAAATGGGAAAGTGACTATACGGGTAACTTAACGATTAATGAAAGGCCCAGTGCACGATGGGGAAGCTGGATCACTATAACGGTCAATCAGGACGTTATTTTCCAGACTTTTTTATTTCCGTTGAAAAGAGACTTCGAGAAAACTGTCGTCTTTGCACTGATTCAAACTGAAGAAGCACTAAATCGTCGCCAGATAAATCAGGCGTTATTAAGTACGGGCGATTTGGCGCATGATGAATTCTAA |
| CsgF |  | ATGCGTGTCAAACATGCAGTAGTTCTACTCATGCTTATTTCGCCATTAAGTTGGGCTGGAACCATGACTTTCCAGTTCCGTAATCCAAACTTTGGTGGTAACCCAAATAATGGCGCTTTTTTATTAAATAGCGCTCAGGCCCAAAACTCTTATAAAGATCCGAGCTATAACGATGACTTTGGTATTGAAACACCCTCAGCGTTAGATAACTTTACTCAGGCCATCCAGTCACAAATTTTAGGTGGGCTACTGTCGAATATTAATACCGGTAAACCGGGCCGCATGGTGACCAACGATTATATTGTCGATATTGCCAACCGCGATGGTCAATTGCAGTTGAACGTGACAGATCGTAAAACCGGACAAACCTCGACCATCCAGGTTTCGGGTTTACAAAATAACTCAACCGATTTTTAA |
| CsgG |  | ATGCAGCGCTTATTTCTTTTGGTTGCCGTCATGTTACTGAGCGGATGCTTAACCGCCCCGCCTAAAGAAGCCGCCAGACCGACATTAATGCCTCGTGCTCAGAGCTACAAAGATTTGACCCATCTGCCAGCGCCGACGGGTAAAATCTTTGTTTCGGTATACAACATTCAGGACGAAACCGGGCAATTTAAACCCTACCCGGCAAGTAACTTCTCCACTGCTGTTCCGCAAAGCGCCACGGCAATGCTGGTCACGGCACTGAAAGATTCTCGCTGGTTTATACCGCTGGAGCGCCAGGGCTTACAAAACCTGCTTAACGAGCGCAAGATTATTCGTGCGGCACAAGAAAACGGCACGGTTGCCATTAATAACCGAATCCCGCTGCAATCTTTAACGGCGGCAAATATCATGGTTGAAGGTTCGATTATCGGTTATGAAAGCAACGTCAAATCTGGCGGGGTTGGGGCAAGATATTTTGGCATCGGTGCCGACACGCAATACCAGCTCGATCAGATTGCCGTGAACCTGCGCGTCGTCAATGTGAGTACCGGCGAGATCCTTTCTTCGGTGAACACCAGTAAGACGATACTTTCCTATGAAGTTCAGGCCGGGGTTTTCCGCTTTATTGACTACCAGCGCTTGCTTGAAGGGGAAGTGGGTTACACCTCGAACGAACCTGTTATGCTGTGCCTGATGTCGGCTATCGAAACAGGGGTCATTTTCCTGATTAATGATGGTATCGACCGTGGTCTGTGGGATTTGCAAAATAAAGCAGAACGGCAGAATGACATTCTGGTGAAATACCGCCATATGTCGGTTCCACCGGAATCCTGA |
| CP19K | Barnacle cement protein with 19 kDa molecular weight | GTTCCGCCGCCGTGCGATCTGAGCATCAAATCTAAACTGAAACAGGTTGGTGCTACCGCGGGTAACGCGGCGGTTACCACCACCGGTACCACCTCTGGTAGCGGCGTTGTTAAATGTGTTGTTCGTACCCCGACCTCTGTTGAAAAGAAAGCGGCTGTTGGTAACACCGGTCTGTCTGCGGTTAGCGCGTCTGCGGCGAACGGTTTCTTCAAAAACCTGGGTAAAGCGACCACCGAAGTTAAAACCACCAAAGATGGTACCAAAGTTAAAACCAAAACCGCGGGTAAAGGTAAAACCGGTGGTACCGCGACCACCATCCAGATCGCTGATGCTAACGGCGGTGTTTCTGAAAAATCTCTGAAACTGGATCTGCTGACCGATGGCCTGAAATTCGTTAAAGTTACCGAAAAGAAACAGGGTACCGCTACCTCTTCTTCTGGTCATAAAGCTAGCGGTGTTGGTCATTCTGTTTTCAAAGTTCTGAACGAAGCTGAAACCGAACTGGAACTGAAAGGTCTGCACCACCACCACCACCACTAA |
| CP43K | Barnacle cement protein with 43 kDa molecular weight | AGTATCAAGCCGCCTGCCGCACTGACTCGCCGCAGTGCCGCGCACCCCTCCGCGGATGCGCCATCTGACCCAACTATGCTGCCAGCGGCCATCCTTCTTCTGAGCTTGGGTGCTGCTTTATCAGCTCCTGCTCCGGGCGTAACACCGCCGGTATCTCCACCCTTACCCCCAGTACCTCCGCCGCTTCCCCCTAAGCGCGCAGCCACCGACGCCGACGCGGTAACTGTCGGTACCTTGAAGACTGCCGGAACCGCCATCGGTAAGTCTTCGGGCGGTGCCGTATCGTTGGAACAGGGCTCGTCGACTGTCTCAAATGCAAACTCAAAGACCGGCGGCACGTCATCGGGGTCCGCCGGTACGGACGCAGCAGGGAAAGCCTCTAGTCGCGGGATTGGAGATGGAACAACCTCACGTGCGGACTCCCAAACAAAAACGAGTACAACTGGCGACGGACGCTCCGAGGCCGACCAACGTTCCACGGGGACAGGTACCACCGGACGTAAGCGTGGCGCGCTGGGCGCCGAAACATCTGCACAGACAACTGGCTCATCGGCGACTGTCGGAGGTGGCAGCGACTCTAAGGGCGAATCTTCTGCGGGGGGGACCGCGAATCAAGGTTCGAATGTTGCCGCTGAATCAGATAGTAATCAGAAGATTCGCAGCACCCGCACGGGCAGTTCGGCGGTTGATGCGAAATCAGGGTCCGCAGCGGCATTGGGGGCCATTAAAGACAAATTGGTTGGAAAATCGGACGCAGCATCTGGGGGGAGTGCAGAGAGCGTGGGCTCGGCGAAAACTGATTTCAACACCGGAGGGTCAGCGGGGCATTCTGCAGGGGAAGGTTCTGGGTTTGCAGAAACGAGTGTCGGCGGGCAGACTCGCCAAACGGGAGCGGTCGAGGGATCACAAACCAGTTCTGCCTCAGGGTCCGTGACCCTTAAGCGTCCTGTGTGGCCTTGTCGCTTACCGTCCAAAGCGCCGAAGGACTGGTTACACGGCTGGGTCCCCGGCACTAAACTTGTGTGGCACTGTGTGTTCCCACATAAGATTCCGGCAAAGTACAGTCAGCTGTACAAGCCTAAGTGGCACCACCACCACCACCACTGA |
| TFF3 | A small secretory peptide from the trefoil factor family | GGCGGCGGTTCTCACCACCACCATCACCACGGTGGTGGTTCTGAAGAATACGTAGGTCTGTCCGCTAACCAGTGCGCTGTACCGGCTAAAGACCGTGTTGATTGTGGCTATCCGCACGTGACGCCGAAAGAATGTAACAACCGTGGTTGCTGCTTCGATTCTCGTATTCCAGGTGTTCCGTGGTGTTTCAAACCGCTGCAAGAAGCGGAATGCACCTTCTAA |
| SEC+N22 | Native CsgA secretion peptide | ATGAAACTTTTAAAAGTAGCAGCAATTGCAGCAATCGTATTCTCCGGTAGCGCTCTGGCAGGTGTTGTTCCTCAGTACGGCGGCGGCGGTAACCACGGTGGTGGCGGTAATAATAGCGGCCCAAAT |

**Table S4**. **Plasmids used in this study.** Links to annotated plasmid sequences are provided for all constructs.

| Plasmid | Description | Source |
| --- | --- | --- |
| [Heme-luciferase](https://benchling.com/s/seq-t06Pb1i81DrZOU45pK3t?m=slm-sO9iImPgC63xj9Q44WzQ) | Prototype plasmid encoded the components of a blood-inducible circuit, including ChuA, HrtR, and a luciferase reporter cassette. The luciferase expression was regulated by the P*_L(HrtO)_* promoter, which was activated in the presence of blood/heme. | Mimee et al^4^. |
| [Heme-sfGFP](https://benchling.com/s/seq-11bP4nFe27H4fzjZOHwB?m=slm-aji9QV0fCJs3XGDmTW1b) | Modified version of the Heme-luciferase plasmid, with luciferase replaced by sfGFP for simpler characterization. | This study |
| [pXW109 Hg(RS-RinA-E11)2](https://benchling.com/s/seq-h52hrUZYc8oAEIPdKW5r?m=slm-IVZGQ8XtzNjQxSxrsLs5) | An optimized biosensing plasmid designed for toxic metal detection featuring a three-layered transcriptional amplifier system that sequentially enhanced output expression levels. | Wan et al^5^. |
| [Sensing plasmid](https://benchling.com/s/seq-UFrwru8P60I7WbHznE1D?m=slm-EiqUb6RCVvAsOaij1NPP) | A pSC101-origin plasmid that included a three-layered transcriptional amplifier regulated by the optimized blood-responsive promoter P*_L_*_(_*_6× HrtO_*_)._ | This study |
| [sfGFP output plasmid](https://benchling.com/s/seq-JHsSD7HabR3JgoEZRp16?m=slm-4X7D0HwvohKJuvWXjAsc) | ColE-origin plasmid with constitutively expressed ChuA and HrtR proteins, paired with the amplifier-specific promoter P*_ecf11_*, which regulated sfGFP expression as a reporter. | This study |
| [CP19K L-glue output plasmid](https://benchling.com/s/seq-0pzb1afN5ETV4ivX4Y8G?m=slm-renaP2p4elR6yJqi3iDL) | ColE-origin plasmid that constitutively expressed ChuA and HrtR, with the amplifier-specific promoter P*_ecf11_* regulating the expression of the barnacle cement protein CP19K and the curli secretion-associated proteins CsgE, CsgF, and CsgG. This modification allowed the engineered strain to secrete the CP19K-based glue matrix upon induction with blood. | This study |
| [CP43K L-glue output plasmid](https://benchling.com/s/seq-0eJHW9kJtjQkAys1y7sF?m=slm-lBI2xJifYKFtHS9vCeGE) | Modified version of the CP19K L-glue plasmid, where CP19K was replaced by CP43K*,* another barnacle cement protein. This modification allowed the secretion of a CP43K-based glue matrix upon blood induction. | This study |
| [TFF3 output plasmid](https://benchling.com/s/seq-7C8HHRCxX27z9HE2Gqxj?m=slm-kvZ73TQh32RqmFY1kxo4) | Modified version of the CP19K L-glue plasmid, with CP19K replaced by *TFF3*, enabling secretion of TFF3 upon blood. | This study |
| [TL-glue output plasmid](https://benchling.com/s/seq-z5HHGPtdlQ6IWp5mZzcn?m=slm-3opOOqvYnOTM2GqzZnjO) | Variant of the CP43K L-glue plasmid, modified to insert TFF3 downstream of CP43K, separated by an *RBS* sequence. This design enabled the engineered strain to secrete both the CP43K-based glue matrix and TFF3 peptide upon blood induction. | This study |
| [miRFP 680-hemeoxygenase](https://benchling.com/s/seq-esSXWZyoOXW40edvGL6a?m=slm-lsfcTV4Nwkz25KUEjGay) | ColE-origin plasmid containing constitutively expressed ChuA and HrtR proteins, with the amplifier-specific promoter P*_ecf11_* regulating the miRFP680 as a reporter for fluorescence imaging. | This study |

**Table S5.** **qPCR primers used in this study.**

| Target genes |  | Primers sequences |
| --- | --- | --- |
| TNFα | Forward | CCTCTCTCTAATCAGCCCTCTG |
| TNFα | Reverse | GAGGACCTGGGAGTAGATGAG |
| IL-6 | Forward | ACTCACCTCTTCAGAACGAATTG |
| IL-6 | Reverse | CCATCTTTGGAAGGTTCAGGTTG |
| COX-2 | Forward | ACAGATGCAATTCCCGGACGTCTA |
| COX-2 | Reverse | GGCATGAAACTGTGGTTTGCTCCA |
| iNOS | Forward | TTCAGTATCACAACCTCAGCAAG |
| iNOS | Reverse | TGGACCTGCAAGTTAAAATCCC |
| hGAPDH | Forward | CTGGGCTACACTGAGCACC |
| hGAPDH | Reverse | AAGTGGTCGTTGAGGGCAATG |
